# Molecular basis for CSB stimulation of the SNM1A DNA repair nuclease

**DOI:** 10.1101/2024.09.05.611390

**Authors:** Rebecca Roddan, Matthew A. Schaich, William J. Nathan, Iulia Kis, Andrea Pierangelini, Lucy R. Henderson, Raman van Wee, Louise Martin, Afaf El-Sagheer, Philipp Kukura, Tom Brown, Darragh O’Brien, Joseph A. Newman, David M. Wilson, Christopher J. Schofield, Bennett Van Houten, Peter J. McHugh

**Affiliations:** Department of Oncology, MRC Weatherall Institute of Molecular Medicine, University of Oxford, Oxford, OX3 9DS, U.K; Department of Pharmacology and Chemical Biology, University of Pittsburgh School of Medicine, Pittsburgh, PA, USA; UPMC-Hillman Cancer Center, Pittsburgh, PA, 15232, USA; Laboratory of Genome Integrity, National Cancer Institute, NIH, Bethesda, MD, USA; Target Discovery Institute, Centre for Medicines Discovery, Nuffield Department of Medicine, University of Oxford, Oxford, OX3 7FZ, U.K; Physical and Theoretical Chemistry, Department of Chemistry, Kavli Institute for Nanoscience Discovery, Dorothy Crowfoot Hodgkin Building, University of Oxford, South Parks Road, Oxford OX1 3QU, U.K; Department of Physiology, Anatomy and Genetics, University of Oxford, OX1 3QX, U.K; School of Chemistry, University of Southampton, SO17 1BJ, U.K; Chemistry Research Laboratory, 12 Mansfield Road, Department of Chemistry, University of Oxford, Oxford, OX1 3TA, U.K; Centre for Medicines Discovery, University of Oxford, NDMRB, Oxford, OX3 7DQ, U.K; Biomedical Research Institute, Hasselt University, 3590 Diepenbeek, Belgium; Ineos Oxford Institute for Antimicrobial Research, University of Oxford, Oxford, OX1 3TA, U.K

## Abstract

The Cockayne Syndrome B (CSB, *ERCC6*) protein, interacts with the exonuclease SNM1A during transcription-coupled DNA interstrand (ICL) repair, with CSB facilitating localisation of SNM1A to ICL damage. The functional and mechanistic details of this interaction in DNA repair, however, have not been defined. Here, we demonstrate that CSB enhances SNM1A resection through ICLs and identify a specific interaction between the winged-helix domain of CSB and the nuclease core of SNM1A that is crucial for recruitment and enhancement of nuclease degradation. Biochemical and single-molecule studies on DNA containing site-specific ICLs reveal that CSB increases the affinity of SNM1A to damaged DNA substrates and also alters the substrate conformation to enhance ICL processing by SNM1A. Notably, CSB was observed preferentially as a dimer when colocalised with SNM1A at ICLs, constrasting with its monomeric nature observed during repair initiation in classical transcription-coupled nucleotide excision repair. The combined results provide molecular insights into the basis of a direct contribution of CSB to a DNA repair reaction.

## Introduction

The integrity of cellular DNA is under constant threat from genotoxic agents, which modify its structure and thus interfere with fundamental cellular processes, in particular DNA replication and transcription. Chemicals that covalently link the two strands of the DNA duplex produce interstrand crosslinks (ICL), lesions that prevent strand separation and are thus particularly problematic^1^. ICL induction via exogenous agents is a widely used cytotoxic chemotherapeutic strategy that targets mainly rapidly dividing cells^2^. ICLs can also be formed by endogenous aldehydes produced during normal cellular metabolism or *via* abasic sites created spontaneously or during DNA repair^3,4^. To remove ICL damage, efficient DNA repair pathways have evolved, employing different factors, dependent on the genomic context and cell cycle phase including; global-genome repair, transcription-coupled (TC) repair (both of which are theoretically operational throughout the cell cycle), and replication-coupled repair (acting in S-phase)^1^.

To date, studies on mammalian ICL repair mechanisms have primarily focused on replication-coupled ICL repair by the Fanconi anaemia (FA) pathway, in which the key ICL ‘unhooking’ incision step is performed by the heterodimeric, structure-selective endonuclease, XPF-ERCC1^5^. After ICL unhooking, XPF-ERCC1 is proposed to collaborate with the 5′-to-3′ exonuclease SNM1A, a metallo-β-lactamase (MBL) fold protein, to further process the ICL, there is also evidence that alternative nucleases including FAN1 are involved in ICL processing^6^. SNM1A-deficient cells exhibit increased cell killing due to ICLs that is epistatic with XPF-ERCC1 loss^7^ and *in vitro* an incision by XPF-ERCC1 results in a nick to the DNA backbone, forming a 5′ phosphate group that presents a loading site for SNM1A exonuclease activity. SNM1A can then processively hydrolyse the phosphodiester backbone of the DNA, and uniquely, digest past the ICL in the 5′-3′ direction^7^. Downstream repair is performed by translesion synthesis and homologous recombination. Although most cells in the body are non-dividing^8^, it is unclear how ICL repair occurs outside of S-phase. Mechanistic insights on replication-independent ICL repair are crucial to understand how ICL-inducing agents impact mostly non-dividing tissues, and this extends to stem-like cancer cells that are regarded as chemoresistant^9^. There is emerging evidence for a role of SNM1A in ICL repair outside S-phase. Most notably, SNM1A (*DCLRE1A*) has been shown to directly interact with the TC-DNA repair factor, Cockayne Syndrome Protein B (CSB)^10^. CSB (*ERCC6*) is a large (168 kDa), SWI/SNF family ATPase^11,12^ linked to a rare segmental progeria, termed Cockayne Syndrome (CS) which features neurodevelopmental defects/neurodegeneration, growth failure, photosensitivity, and a premature ageing phenotype^13^. CSB has a fundamental role in recognising DNA damage-stalled RNA Polymerase II (RNAPII) during transcription, with CS phenotypes proposed to be caused by defects in DNA repair and transcription recovery^14^. The CSB-RNAPII recognition event initiates recruitment of downstream DNA repair factors that act to remove and replace the damaged DNA. The role of CSB in TC repair has been extensively studied in the context of UV irradiation-induced damage (*via* NER), although there is emerging evidence for CSB guiding resolution of other types of DNA damage including DNA-protein crosslinks, oxidative DNA damage and ICLs^10,15–22^.

A potential physical interaction between CSB and SNM1A was initially identified following a yeast two-hybrid screen^10^. Purified recombinant SNM1A and CSB were also shown to interact in cells, with CSB enhancing the exonuclease activity of SNM1A on simple single-stranded (ss) and double-stranded (ds) DNA oligonucleotides^10^. Fluorescently tagged CSB and SNM1A were shown by confocal microscopy, to co-localise at sites of ICLs in a transcription-dependent manner^10^. CSB deficient cells exhibit sensitivity to ICL-inducing agents, particularly in G_0_/G_1_ where replication-coupled pathways are not available^10^. CSB-deficiency resulted in reduced ICL ‘unhooking’ in non-cycling, differentiated neural cells, where there was an association with persistence of γ-H2AX DNA damage foci. Further yeast two-hybrid studies suggested that the C-terminal domain of CSB (amino acids 1187-1493) interacts with the catalytic domain of SNM1A (amino acids 698-1040), thus narrowing down the interacting regions. Overall, these studies led to the proposal that CSB co-ordinates ICL resolution through SNM1A in a transcription-dependent manner^10^. However, the molecular interactions governing this process, however, have not been defined.

Our work here addresses the molecular basis of CSB stimulation of SNM1A nuclease activity. We describe the structural and molecular-level requirements for CSB to interact with SNM1A, defining the nature of the interaction between these two proteins and relevant DNA substrates. We additionally identify a distinct intermolecular interaction between CSB and SNM1A that is crucial for their cellular interaction and SNM1A recruitment. We show that CSB stimulates SNM1A nuclease activity during biochemically reconstituted ICL unhooking reactions, in a manner unrelated to CSB’s translocase/helicase activity and only partly dependent upon direct interaction between the two proteins. Single-molecule studies with fluorescently-labeled nuclear extracts reveal the oligomerization state of CSB on damaged and undamaged DNA, and crucially, identify that CSB recruits SNM1A to ICL damage, substantially increasing the affinity and lifetime of SNM1A on target substrates. Together, our work provides key molecular insights into the coordinated damage removal process involving CSB and SNM1A. Finally, this work suggests a working model of how TC dependent ICL repair is achieved.

## Results

### The CSB:SNM1A interaction requires the winged-helix domain of CSB

Prior work implies that CSB and SNM1A likely interact via their respective C-terminal regions^10^. We initially sought to replicate this finding biochemically and investigate in detail how the two proteins interact in the presence of other repair factors. CSB protein (168 kDa) contains a central, folded ATPase domain (amino acids (a.a.) 402-1040) with high sequence homology to SWI/SNF chromatin remodelers^23^. Flanking the ATPase domain of CSB are extended N- and C-terminal domains about which little is known from a structural perspective. The N-terminal domain (a.a. 1-401) of CSB has been shown to have an N-terminal coiled coil motif, but is otherwise predicted to be largely disordered (a simple domain map of CSB in Fig. 1A). The C-terminal CSB domain (a.a. 1187-1493) is also mostly disordered, with the exception of a ubiquitin-binding and winged-helix domain (WHD, a.a 1424-1493) ^24^.

The C-terminal region of SNM1A (a.a. 698-1040) and full-length CSB were purified from insect cells, with representative C-terminal constructs of CSB (a.a. 1187-1493, 1187-1308, 1187-1401, 1308-1493, 1424-1493) expressed in *E. coli* (Fig. S1). The core catalytic, C-terminal domain of SNM1A (a.a. 698-1040, containing the MBL-β-CASP domains), was used rather than full-length protein due to established stability issues^10^ – hereafter this is referred to as SNM1A. This form of SNM1A has been previously shown to have the same exonuclease activity profile as the full-length version^10^. Surface plasmon resonance (SPR) was used to define the affinities of CSB constructs to immobilized N-Avi-Biotin-SNM1A. The high affinity interaction between the C-terminal and full-length CSB constructs and SNM1A were comparable (apparent dissociation constants (K_D_s) of 10.7 vs. 4.8 nM, Fig. 1B). No SNM1A binding was observed with the C-terminal region of CSB (a.a. 1187-1401) lacking the WHD of CSB (a.a. 1424-1493) (Figure 2A). This observation indicates that the WHD is the minimal interaction motif for binding to SNM1A, while, the K_D_ of 89 nM (CSB-WHD:SNM1A) indicates that optimal binding is only conferred by the entire CSB C-terminal region. Dose-response curves were also obtained for immobilised SNM1A with CSB_1187-1493_, CSB_1308-1493_ and CSB_1424-1493_ by multicycle SPR analysis (Fig. S2) - apparent K_D_ values obtained were of the same trend as single cycle values (Fig. 1B).

We next sought to define the molecular contacts governing the SNM1A:CSB interaction. AlphaFold3 (AF3) modelling was performed with C-terminal SNM1A (a.a. 698-1040) and full-length CSB (Fig 1C). Statistical analysis of the modelled structure gave an overall iPTM of 0.81 and pTM of 0.37, suggesting high confidence in the predicted interaction. No secondary structure was predicted for large regions of CSB (a.a. 1-76, 296-412 and 1025-1250). Hydrogen bonding interactions are predicted between SNM1A and the CSB-WHD (SNM1A-E864:CSB-T1447, R1478 and SNM1A-L198:CSB-R1467). Additionally, interactions were observed with the extended CSB C-terminal domain (SNM1A-E915:CSB-S1329,R1330) and the ATPase domain, lobe 1 (SNM1A-R824:CSB-D634).

To experimentally validate the AF3 model and understand whether SNM1A binding induces additional structural changes in the CSB protein, hydrogen-deuterium exchange mass-spectrometry (HDX-MS) experiments were performed where good sequence coverage (∼80%) was achieved for the full-length protein (Fig 1D and S3-5). Strikingly, the only region of the CSB protein that exhibited altered and decreased deuterium uptake upon SNM1A binding was the CSB WHD. After 30 s of incubation, decreased deuterium uptake was observed in a.a. 1439-1453 and 1478-1487, with decreased uptake in additional regions of the WHD, a.a. 1428-1437 and 1467-1476 being observed after 10 min (Fig 1D). Even though we obtained peptide coverage within the region a.a. 1460-1466, no changes in deuterium uptake were observed. Considering no changes in deuterium uptake were observed throughout the rest of the full-length CSB protein, this indicates that SNM1A binding forms an interface with the majority of the WHD, inducing conformational changes within the WHD itself.

Together, our data demonstrates that the WHD of CSB is the minimal interaction motif for interaction with SNM1A. Nonetheless, our SPR data show that the remaining CSB C-terminus (a.a. 1187-1423) is required for high affinity SNM1A binding, perhaps by stabilising or orientating the WHD conformation for optimal SNM1A binding.

**Figure 1:**
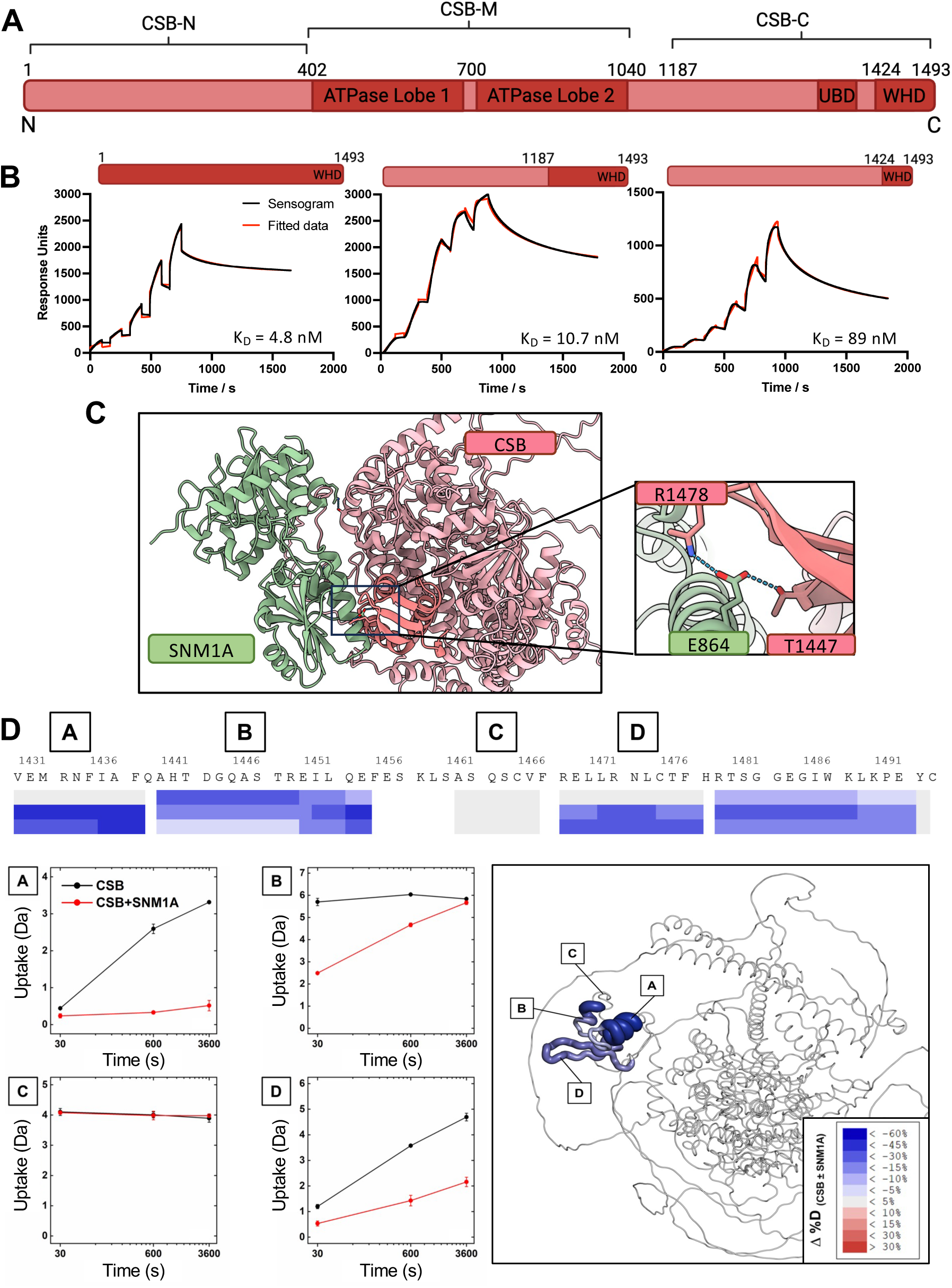
Defining the molecular interaction between CSB and SNM1A. **A.** Domain map of CSB, defining the N-terminal (CSB-N), ATPase (CSB-M) and C-terminal (CSB-C) domains. **B.** Single-cycle response curves from SPR analyses, with apparent equilibrium dissociation constants given. Parameters determined from fitting are given in Table S1. **C**. AlphaFold3 modelling of CSB (full-length, a.a. 1-1493, red) and SNM1A (C-terminal domain, a.a. 698-1040, green) with hydrogen bonding interaction between CSB-WHD and SNM1A highlighted. **D.** HDX-MS of CSB interaction with SNM1A. Differential deuterium uptake was observed in the WHD domain of CSB in the presence SNM1A, exemplified by peptides A, B, and D. No differences between apo- and holo-states were observed for the majority of the protein, an example of which is represented by peptide C. HDX-MS changes were mapped to the AlphaFold model of full-length CSB

### Identification of CSB residues required for specific interaction with SNM1A

The key molecular contact identified by AF3 modelling was SNM1A-E864:CSB-T1447, well supported by HDX-MS analysis (within the region a.a. 1439-1453). This interaction site was probed further using CSB and SNM1A constructs with alanine substitutions at these two residues. The interaction between wild-type (WT) versus E864A-SNM1A (MBL-β-CASP domain, a.a. 698-1040) and full-length CSB was found to be of comparable affinity (K_D_ = 4 vs. 8 nM) by SPR (Fig 2A, Table S2), while interactions of E864A-SNM1A with the CSB-WHD and CSB C-terminal domains were reduced (K_D_ = 89 nM vs. *ca.* 1 μM and K_D_ = 12 vs. 29 nM, Fig S1), consistent with the AF3 modelling. Data obtained with CSB-T1447A was more striking; no interaction was observed between WT-SNM1A and full-length or WHD-CSB with T1447A substitution (Fig 2B, 2C and S6). Circular dichroism (CD) measurements of the CSB-WHD (comparing WT and T1447A CSB) confirmed that this observation was due to the alanine substitution rather than misfolding of the protein domain (Fig. S7). These results are also consistent with our HDX-MS results, where CSB-T1447 is present in the region of CSB that is involved in the initial interaction with SNM1A (a.a. 1439-1453).

We investigated the presence of biophysically defined CSB-SNM1A interactions in cells. Co-immunoprecipitation experiments were performed using transiently expressed GFP-tagged SNM1A or CSB constructs with the relevant alanine substitutions (Fig 2D and E). CSB was co-immunoprecipitated by both WT and a nuclease-deficient variant of SNM1A, D736A. Confirming the importance of the E864-SNM1A residue in CSB binding, a significantly diminished interaction with CSB was observed with E864A-SNM1A relative to WT SNM1A (Fig. 2D). Consistent with our SPR findings, T1447A-CSB did not interact with SNM1A in cells (Fig. 2E). Importantly, the alanine substitution at T1447-CSB was not observed to alter the interaction with the major subunit of RNAPII, RPB1.

**Figure 2:**
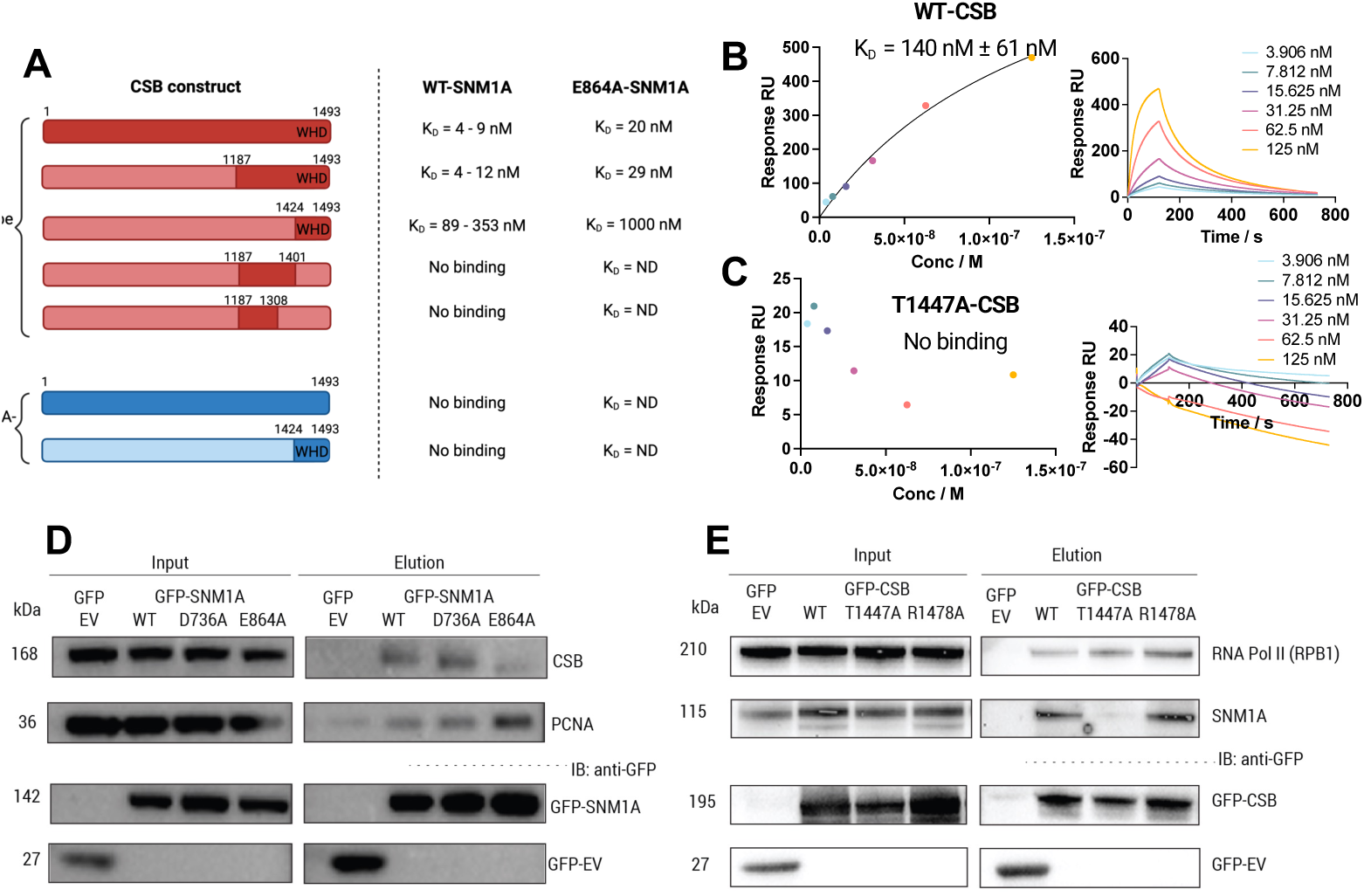
Identification of a CSB:SNM1A separation-of function variant. A. Apparent equilibrium dissociation constants obtained between different CSB (vary a.a. lengths, wild-type and T1447A) and SNM1A (a.a. 698-1040) constructs. The dark red/blue regions of the protein sequence maps represent the amino acid ranges used for experiments. ND = Not determined. B. Multi-cycle SPR analyses of immobilized N-His-ZB-CSB-Avi-Biotin (WT) with SNM1A (WT, a.a. 698-1040, untagged) as the analyte. C. Multi-cycle SPR analyses of immobilized N-His-ZB-CSB-Avi-Biotin (T1447A) with SNM1A (WT, a.a. 698-1040, untagged) as the analyte. D. Co-immunoprecipitation experiment with immobilised GFP or GFP-tagged SNM1A (full-length construct: WT, nuclease-dead variant D736A or E864A). **E.** Co-immunoprecipitation experiment with immobilised GFP or GFP-tagged CSB (full-length construct: WT, T1447A or R1478A).

### The molecular mechanism by which CSB facilitates SNM1A-dependent DNA damage processing

CSB enhancement of SNM1A exonuclease activity has previously been observed on simple single-stranded (ss) and double-stranded (ds) DNA substrates^28^. Ee expanded the substrate range and examined SNM1A exonuclease activity in the presence CSB across a panel of potentially relevant oligonucledotide DNA repair intermediate substrates (Table S3 and Fig. S8). All substrates were designed with a phosphate at the 5′ end of the 3′-^32^P-labelled DNA strand (5′P), a required feature for SNM1A nuclease activity, while the 5′ end of the strand not under analysis was capped with a biotin group (Bio) to prevent competing digestion. As expected^29^, SNM1A alone has a clear preference for ssDNA, with significantly reduced hydrolysis observed on dsDNA (27% vs. 79% digestion on dsDNA vs. ssDNA after 10 min for SNM1A alone). On a substrates with a recessed 5’ end (3’ overhang substrate), or containing a 5’ overhang substrate, SNM1A can also be observed slowing when entering the region of dsDNA (Fig. S9). Full-length CSB enhanced the nuclease activity of SNM1A on all substrates, albeit with only a minor effect on the 5’ overhang substrate. SNM1A alone has poor activity on the 17 bp bubble DNA substrate, which models a transcriptional intermediate (83% substrate remaining after 10 min), but in the presence of CSB, a ∼5-fold stimulation was observed with only 12% initial substrate remaining at 10 min.

We sought to build on these insights to understand the biochemical basis of the observation that CSB is required for efficient ICL unhooking in non-dividing cells^28^. To specifically and quantitatively assess whether CSB has a role in stimulating SNM1A resection past an ICL to achieve unhooking, a substrate (herein referred to as the ‘0-nucleotide’ substrate) was designed whereby SNM1A encounters a triazole DNA interstrand crosslink in its first hydrolytic event, i.e., there are no nucleotides preceding (5’ to) the crosslink (Fig. S10). This ICL produces relatively minor helical distortion^30,31^, similar to several major classes of physiologically important ICLs^32^. Hydrolysis of the substrate by SNM1A alone was partial, with ∼40% of the substrate digested after 5 min (Fig. 3A and 3B). With the addition of CSB, only ∼15% of the substrate remained, suggesting that CSB has a substantial role in promoting SNM1A digestion through ICLs^43^.

We next investigated the basis of increased apparent SNM1A nuclease activity in the presence of CSB. We hypothesised that this could be attributed to one, or a combination, of the following factors: 1) enhanced SNM1A recruitment to substrate, 2) substrate structure modulation, 3) a direct physical interaction that enhances SNM1A catalysis through increased processivity, or 4) increased turnover by stimulating product release. Nuclease assays were performed on the 0-nucleotide ICL substrate comparing SNM1A activities in the presence of WT-CSB or T1447A-CSB, i.e., where CSB and SNM1A do and do not physically interact, respectively (Fig. 3A and 3B). A degree of stimulation of SNM1A activity was observed with T1447A-CSB, although this was reduced (38% vs 63% substrate remaining after 10 min for WT vs. T1447A-CSB). We conclude that while the physical interaction between CSB and SNM1A is important for SNM1A stimulation, other features of CSB contribute. To assess whether the additional stimulation with WT-CSB was associated with substrate modulation^44^, we performed the same assay on ssDNA (Fig. 3C and 3D), a flexible unstructured molecule lacking the complex duplex DNA structural features present in the ICL repair intermediate. We observed that unlike WT-CSB, T1447A-CSB does not stimulate SNM1A digestion. This observation supports the proposal that both substrate modulation and the specific interaction between CSB and SNM1A are key for stimulation.

Further experiments with E864A-SNM1A confirmed that this alanine substitution had no effect on baseline SNM1A nuclease activity (Fig. 3E and 3F). But by contrast with assays perfomed with T1447A-CSB, no stimulation by wild-type CSB was observed with E864A-SNM1A on the ‘0-nucleotide ICL’ substrate. We hypothesised that contrasting results in the E864A-SNM1A and the T1447A-CSB assays might reflect occlusion of the DNA substrate by the C-terminus of CSB, whereas the T1447-CSB:SNM1A interaction is required to alleviate this and allow SNM1A loading onto the substrate.

We therefore investigated whether the C-terminal domain of CSB is capable of binding to DNA, using electrophoretic mobility shift assay (EMSA). We found that CSB-C can bind a range of DNA repair intermediate-like substrates (Fig 3G and H), and for the first time, observed full-length CSB binding to ssDNA, consistent with its reported ssDNA annealing activity. EMSAs performed with truncated C-terminal constructs showed that the CSB-WHD alone did not exhibit DNA binding (data not shown, although WHDs generally are often DNA binding motifs^33,34^), whereas C-terminal fragments lacking the WHD showed decreased DNA binding relative to full-length protein (Fig. S14).^40^ Strikingly, SNM1A nuclease assays in the presence of individual CSB domains revealed that the presence of the CSB C-terminal domain alone is indeed inhibitory to SNM1A activity (Fig S15). We propose that both a direct protein-protein interaction and DNA substrate modulation are responsible for CSB’s enhancement of SNM1A activity, with CSB aiding loading of SNM1A onto DNA, with significant structural rearrangements of CSB and/or DNA upon CSB:DNA binding.

Finally, we determined whether the ATPase activity of CSB was relevant in this apparent stimulation of SNM1A activity (Fig S11). In the presence of ATP (50,000 molar equivalents to CSB) or with an ATPase-inactive CSB (E647Q mutation, characterisation in Fig S11), no effect on SNM1A stimulation was observed, suggesting that the activity is ATP-independent. (Fig. S11, S12).

**Figure 3:**
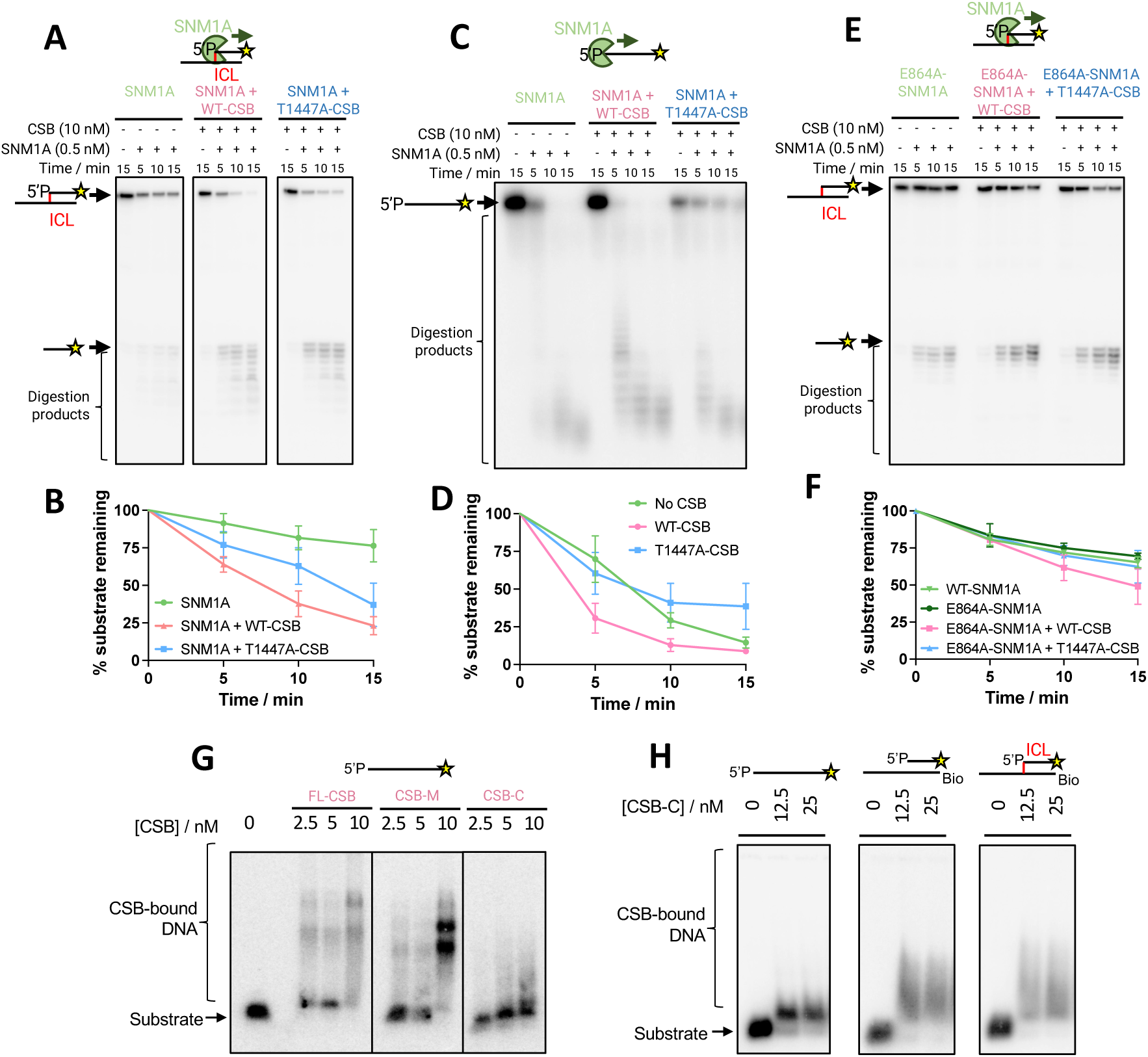
CSB enhances SNM1A digestion through ICLs via SNM1A recruitment and substrate modulation. Oligonucleotides used (with preparation method) are given in Table S3 and Fig. S8. The location of the 3′ ^32^P is given by a yellow star, Bio represents the location of 5’ biotins on the oligo and 5’P the location of a 5’ phosphate group. Quantification was based upon assays conducted in triplicate; error bars represent one standard deviation. **A.** Time course assays of SNM1A activity on the ‘0-nucleotide’ ICL substrate in the absence/presence of CSB (WT or T1447A). **B.** Quantification of the assay shown in B. **C.** Time course assays of SNM1A activity on single-stranded DNA substrate in the absence/presence of CSB (WT or T1447A). **D.** Quantification of the assay shown in panel D. **E.** Time course assays of SNM1A (E864A) activity on single-stranded DNA substrate in the absence/presence of CSB (WT or T1447A). **F**. Quantification of the assay shown in panel F. data for WT-SNM1A is also shown, from data given in panels B and C, as the assays were run in tandem. **G**. EMSA of full-length CSB, CSB ATPase domain (CSB-M, a.a. 402-1014), CSB C-terminal domain (CSB-C) a.a., 1187-1493) binding to single stranded DNA in the presence of 4 mM EDTA. **H**. EMSA of CSB C-terminal domain (a.a. 1187-1493) binding to different DNA structures in the presence of 4 mM EDTA.

### Off-DNA CSB oligomerisation state

Previous structural and biochemical studies of the CSB protein have raised questions regarding its oligomerization state; it has been suggested that CSB is a dimer in solution based on size-exclusion chromatography (SEC) and immunoprecipitation experiments with over-expressed individual CSB domains^35^. However, structural investigations of CSB-containing, TC-NER initiation complexes indicate the presence of only one copy of CSB. It is unclear whether this is due to a lack of structural resolution (due to high flexibility of its numerous unstructured domains) or is functionally relevant^36–39^. Dimerisation is also possibly concentration dependent and/or could be DNA-binding mediated.^35,40^

Consistent with previous work, purification of full-length CSB by SEC and comparison of the observed retention volume with the associated calibration curves indicated that the protein was a dimer (Fig S16). Mass photometry (MP) experiments of full-length CSB, however, indicated that the protein was, predominantly, a monomer in solution (Fig 4A). CSB constructs lacking the N-or C-terminal domains (ΔN-CSB, a.a. 402-1493 and ΔC-CSB, a.a. 1-1014, respectively) were also studied - both were also monomeric in solution, with a small amount (∼7%) of dimer observed for ΔN-CSB (Fig. S17). A concentration-dependent dimerisation of ΔN-CSB was also observed by SEC-MALS (Fig. S18). The differences in results obtained by MP compared with SEC-MALS are likely due to concentration differences, with MP experiments performed at 200 nM compared with 5 μM in the SEC-MALS analyses, with the latter likely driving dimerisation.

To investigate whether SNM1A mediates CSB dimerization in solution, we performed both MP and SEC experiments with both proteins present. MP experiments with equimolar ratios of CSB and SNM1A indicated that a 1:1 complex was formed with K_D_ of 499 ± 215 nM (mean ± standard deviation) (Fig 4B). Additionally, consistent with experiments performed with CSB alone at higher concentrations, co-purification of CSB:SNM1A complexes (with full-length or CSB-C (a.a. 1187-1493)) indicated a 2:1 stoichiometry (Fig. 4C-F). Interestingly, the SNM1A:CSB-WHD complex indicated a 1:1 stoichiometry (Fig. S19 and S20), consistent with the HDX-MS results and AF3 modelling suggesting that SNM1A interacts with CSB’s WHD via a specific interaction requiring CSB-T1447. We conclude that isolated CSB can exist as a monomer or a dimer alone, as is the case when in complex with SNM1A, and this equilibrium position is likely concentration and/or crowding dependent. Since the estimated cellular CSB concentration is ∼31 nM, CSB could potentially be induced to dimerise by local concentrations or other enrichments^41^.

### DNA provides a dimerisation interface for CSB

To obtain direct visualization of individual molecular CSB states in real-time, we employed single-molecule analysis of DNA-binding proteins from nuclear extracts (SMADNE), a technique whereby protein-DNA binding events are studied in a ‘near-nuclear’ environment^42^.

HaloTag-CSB was overexpressed in Expi293F cells, nuclear extracts prepared and expression quantified by SDS-PAGE and immunoblot (Fig. S21 and S22). Two colour experiments were initially performed on lambda DNA (i.e. an undamaged, dsDNA substrate) tethered between two beads on the LUMICKS’ C-Trap. CSB extracts were prepared individually with two JF dyes, JF552 (green) and JF635 (red), and mixed. A representative kymograph, showing the binding of the two CSB colours to dsDNA over time, is given in Fig. 4G. CSB exhibited two modes of binding on undamaged DNA, with both short (7.12 s) or long (73.2 s) lifetimes, with an apparent K_D_ of 1.2 nM (Fig. 4H, Table S5). This two-phase binding regime could reflect two possible CSB conformational states upon dsDNA binding, since CSB is known to be structurally-dynamic, undergoing significant structural rearrangements during DNA repair^43,44^. We noted some two colour colocalisation events (example given in Fig. 3G, event at 352-369 s), suggesting that CSB can form dimers while interacting on dsDNA. We also observed events where two CSB colour proteins diffuse together on DNA, which is indicative of colocalisation (Fig. S23). We measured the average photon intensities of the CSB track events, and comparison of the average photon count of all CSB events showed a bimodal distribution, indicative of a combination of monomers and dimers (Fig. S23). Based on this distribution, we propose that average photon counts below the midpoint identify monomeric CSB binding events, and those above, to reveal dimeric events. The difference in track intensities, suggesting monomer or dimer CSB events, can be observed in Fig. 4G. Considering both photon count intensities and two colour colocalisation events, ∼35% of CSB binding events were dimeric.

We studied the mode of molecular assembly of the CSB dimer by examining colocalisation in the two colour experiment. A graphical representation of the possible modes in which the two types of labelled CSB can colocalise is given in Fig. 4I. There are 11 possibilities, where categories 1 and 2 represent each of the two labelled CSB species binding to DNA alone. We found that in 93% of colocalised events (i.e., categories 3-11), categories 3, 5 and 9 were represented, suggesting that one molecule of DNA-bound CSB recruits a second molecule to form a dimer (Fig 4J). Additionally, the lower number of category 6 (representing ∼8% of colocalised events) and absence of category 7 and 8 events suggests that CSB does not substantially dimerise when not bound to DNA at the relatively low (and more physiological) concentrations used in SMADNE. Additionally, by considering photon count intensities of single colour tracks, in 20% of cases, a dimeric CSB bound to DNA. The difference between the two colour colocalisation experiment is likely due to slow exchange of the CSB dimer. These results are consistent with the MP results, where CSB was observed to be monomeric in dilute solution (Fig. 4A). Additionally, in 85% of colocalised events, the second copy of bound CSB is released prior to dissociation of the first bound copy (category 5 or 9). Together, these data demonstrate that DNA helps provide a dimerization interface for CSB.

**Figure 4:**
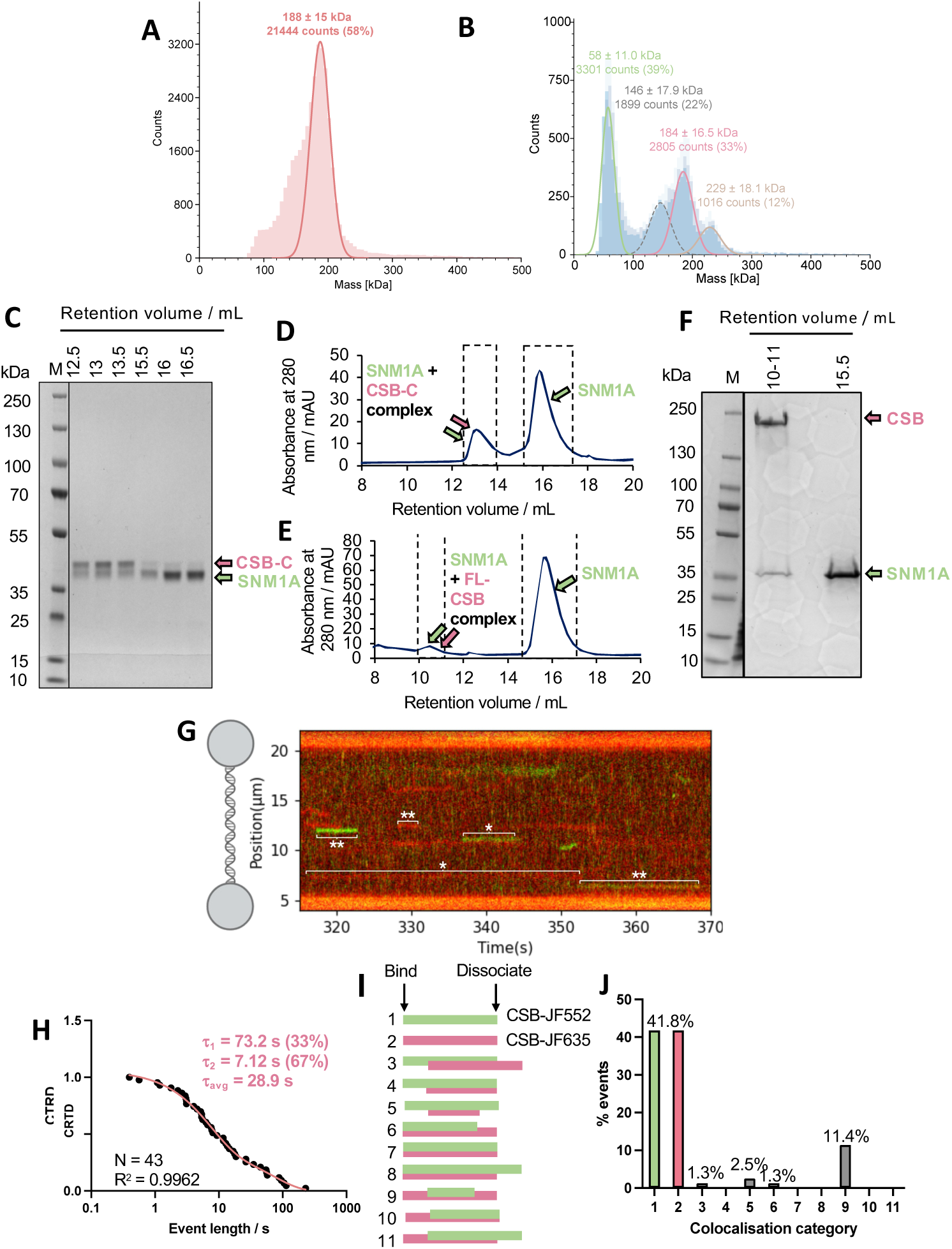
Investigation of the oligomerisation state of CSB. **A.** Mass histograms from mass photometry measurements of the CSB protein. The expected molecular weight of the CSB monomer is 180 kDa. **B.** Mass histogram of experiments of CSB (expected: 180 kDa, observed: 184 kDa) and SNM1A (expected: 36 kDa, observed: 58 kDa) mixed at 200 nM each, indicating that a 1:1 complex of the two proteins forms. Analysis of the mass counts corresponding to each protein state determined a KD of 499 ± 215 nM (mean ± standard deviation) for the interaction between CSB and SNM1A. **C.** Fractions from size-exclusion chromatography from co-purification of CSB-C and N-Avi-Biotin-SNM1A (a.a. 698-1040) analysed by SDS-PAGE. **D.** Size-exclusion chromatography trace from co-purification of CSB-C (a.a. 1187-1493) and SNM1A (a.a. 698-1040). The peak at 12.5-13.5 mL corresponds to a CSB:SNM1A 2:1 complex as supported by SDS-PAGE, MS analysis (for protein identities) and size-exclusion calibration curve from Superdex S200 GL 10/300. See Supplementary Information for calibration curve. **E.** Size-exclusion chromatography trace from co-purification of FL-CSB and SNM1A (a.a. 698-1040). The peak at 10.5 mL corresponds to the CSB:SNM1A 2:1 complex as supported by SDS-PAGE and MS analysis (for protein identification) and quantitative size-exclusion chromatography (Superdex S200 GL 10/300 column). **F.** Fractions from size-exclusion chromatography from co-purification of full-length CSB and SNM1A (a.a. 698-1040) analysed by SDS-PAGE. **G.** Representative kymograph of CSB (2 colour experiment) binding to lambda DNA. **H.** CRTD analysis of CSB binding to lambda DNA. **I.** Schematic of the possible modes of DNA binding for two colour protein experiment with two labelled CSB proteins. **J.** Bar graph showing the representative of different types of molecular assembly events observed with the two colour CSB experiment.

### Real-time insights into CSB:SNM1A interactions on DNA

Following our observations that CSB can potentially dimerise on undamaged dsDNA, we sought to understand the features of the CSB:SNM1A interaction on the same substrate, one in which nuclease stimulation has been observed. Partially-purified HaloTag-JF-552-labelled SNM1A (green, a.a. 698-1040, i.e., the stabilised, catalytic C-terminal domain) was added to the HaloTag-CSB (red JF635-labelled CSB) nuclear extract, as SNM1A overexpression is poorly-tolerated in mammalian cells^45^. In a majority (85%) of CSB:SNM1A colocalisation events (categories 3-11, Fig. 5A-C), a category 9 assembly mechanism occurs, where CSB recruits SNM1A to DNA, with CSB remaining bound after SNM1A is released. Only in very rare cases did CSB and SNM1A land together on the DNA (∼0.2%) (categories 6-8). By studying category 9 events in isolation (utilising the time between the start of CSB binding to the start of SNM1A binding as the association rate), we determined an affinity for SNM1A alone to dsDNA (K_D_ = 22 nM) and to CSB-bound DNA (K_D_ = 18 nM), reflecting an affinity higher than the in-solution affinity between CSB and SNM1A determined by mass photometry (K_D_ = 499 ± 215 nM, Fig 4B and Table S5). These results advance our understanding of CSB-SNM1A coordination by revealing that the preferred mode of binding to DNA is sequential, consistent with our hypothesis that CSB loads SNM1A onto DNA via its WHD.

Analysis of the binding lifetimes of SNM1A alone to dsDNA revealed two distinct lifetimes, likely suggesting that it binds DNA in two different conformations. The DNA binding cleft of SNM1A is highly positively charged, and so there is a possibility of non-productive binding to DNA (i.e., a non-catalytically feasible binding mode) as well as a productive, longer-lived mode, potentially at rare non-specific nicks in DNA that present a 5’P group allowing preferred SNM1A engagement. Comparison of SNM1A binding lifetimes with and without CSB, showed a 2-fold increase in the lifetime in the longer binding regime only (SNM1A alone, τ_1_ = 0.71 s and τ_2_ = 8.6 s and for SNM1A: CSB, τ_1_ = 0.82 and τ_2_ = 16.2 s). This could reflect a specific DNA-bound conformation of CSB that stabilises SNM1A binding. Additionally, we analyzed CSB kymotrack photon counts to determine its oligomerisation state in complex with SNM1A and found that CSB was a dimer in ∼73% of colocalisation events (Table S5). Considering that ∼35% of CSB:DNA events exhibit CSB dimerisation, this suggests a preferential recruitment of SNM1A to the dimer versus the monomer, potentially due to the availability of two WHD binding sites resulting in a higher association rate. Interestingly, comparison of the lifetimes of SNM1A colocalised with monomer or dimer CSB (Fig. 5D and 5E) events suggested that lifetimes were generally 2-fold longer with the monomer rather than the dimer. Overall, this is consistent with a model of CSB regulating SNM1A activity by recruiting and retaining SNM1A on DNA.

**Figure 5:**
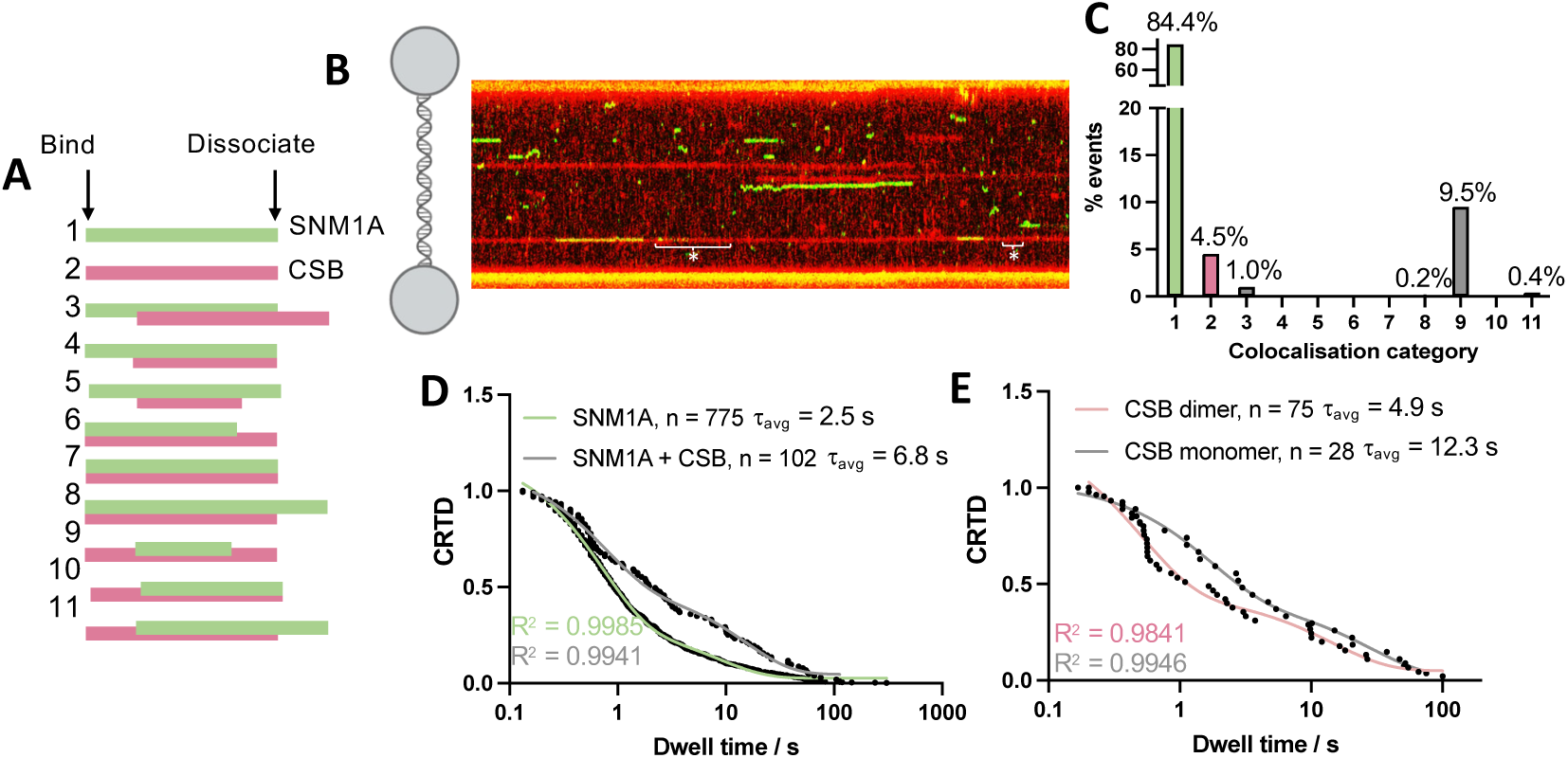
Single-molecule studies of CSB:SNM1A assembly on undamaged DNA. **A.** Graphical representation of the potential modes of molecular assembly of CSB and SNM1A binding and dissociating, alone or together on DNA. **B.** Representative kymograph of CSB (JF-635, red) and SNM1A (JF-552, green) binding to lambda DNA. **C.** Bar graph showing the percentage of events represented by different categories of colocalisation for CSB and SNM1A binding alone or together to lambda DNA. The schematic in panel B is to be used as the key. **D**. CRTD plot of SNM1A dwell times alone (green) or colocalised with CSB in category 9 events (grey) on lambda DNA. Two-phase binding regimes were determined for both types of events, with average binding lifetimes shown here (individual lifetimes of each regime are given in Table S5). **E.** CRTD plot of SNM1A dwell times on monomer (grey) or dimer CSB (pink) on lambda DNA i.e., separating the monomer and dimer CSB in category 9 colocalisation events with SNM1A from panel C. Two-phase binding regimes were fit for each with the average of the two lifetimes given. The lifetime of each binding regime is given in Table S5.

### Dimeric CSB recruits SNM1A to an ICL repair intermediate

Next, we characterised the CSB and SNM1A interaction on a relevant, DNA repair intermediate substrate. We prepared a substrate containing a single site-specific lesion, mimicking a post-incision, ICL-containing, transcription-sized bubble. Briefly, this DNA complex was comprised of a ss flap substrate of 17 bp with a 5’P and an ICL (incorporated via a click reaction as previously described) 3’ to the flap (schematic representation in Fig. 6A, with oligonucleotide sequences and characterisation in Table S4 and Fig. S10). This single, site-specific lesion allows repair-relevant binding and biochemical events of CSB and SNM1A to be studied in isolation. A representative kymograph of CSB (red) and SNM1A (green) binding to this substrate is given in Fig. 6B. CRTD plots of SNM1A binding to the ICL intermediate fit to a one-phase exponential decay (Fig. 6D, green) in contrast to the two-phase decay observed on dsDNA (Fig. 5D), suggesting that SNM1A binds occurs to this damaged substrate in a single conformation. Consistent with colocalisation events observed on dsDNA, CSB:SNM1A colocalisation events on the ICL were predominantly of a mechanism where CSB recruits SNM1A to the damage (70% of events, Fig. 6D, events 9-11 vs. 3-11). Although k_on_ values were comparable (SNM1A: 3.13 ± 3.19 × 10^6^ M^-1^ s^-1^ vs. CSB:SNM1A: 1.87 ± 0.16 × 10^6^ M^-1^ s^-1^, Table S6), a dramatic increase (∼10-fold) in affinity and event lifetimes of SNM1A binding were increased in the presence of CSB (K_D_ = 355 nM vs. 45 nM; τ_avg_ = 0.90 vs. 11.8 s). Longer-lived SNM1A binding could imply that CSB recruits or remodels SNM1A to a more productive, repair-relevant conformation at the repair intermediate. Notably, and in stark contrast, on undamaged lambda dsDNA, the affinity of SNM1A versus SNM1A:CSB was comparable (K_D_ = 21.8 vs. 17.5 nM). Overall, these observations imply that SNM1A binding alone to this repair intermediate is disfavoured, with CSB aiding recruitment and retention of SNM1A at the damage site. Additionally, a striking preference of CSB dimer binding to the ICL repair intermediate was observed (∼78% of events Fig. 6F), and analysis of the photon counts suggested that primarily dimeric CSB bound directly to the DNA, rather than one copy of CSB recruiting another (Fig. S23). This contrasts with CSB binding on dsDNA, where for 24% of DNA binding events, exchange of CSB copies is observed (Fig. 6F). This is also reflected in results that SNM1A preferentially binds dimeric CSB on ICLs (Fig. 6G).

The dimeric oligomerisation state was selective to the ICL repair intermediate, as non-specific binding events to the rest of the tethered DNA were primarily monomeric (i.e., comparable to results observed on lambda DNA. The photon counts are noticeably higher in the centralized CSB binding event (on the ICL) compared with non-specific binding to the rest of the substrate (i.e., ds, lambda DNA) (Fig. 6B).. This observation provides key mechanistic insight into the regulation of SNM1A recruitment to damage and demonstrates that CSB dimerisation is potentially required for TC-ICL processing.

**Figure 6:**
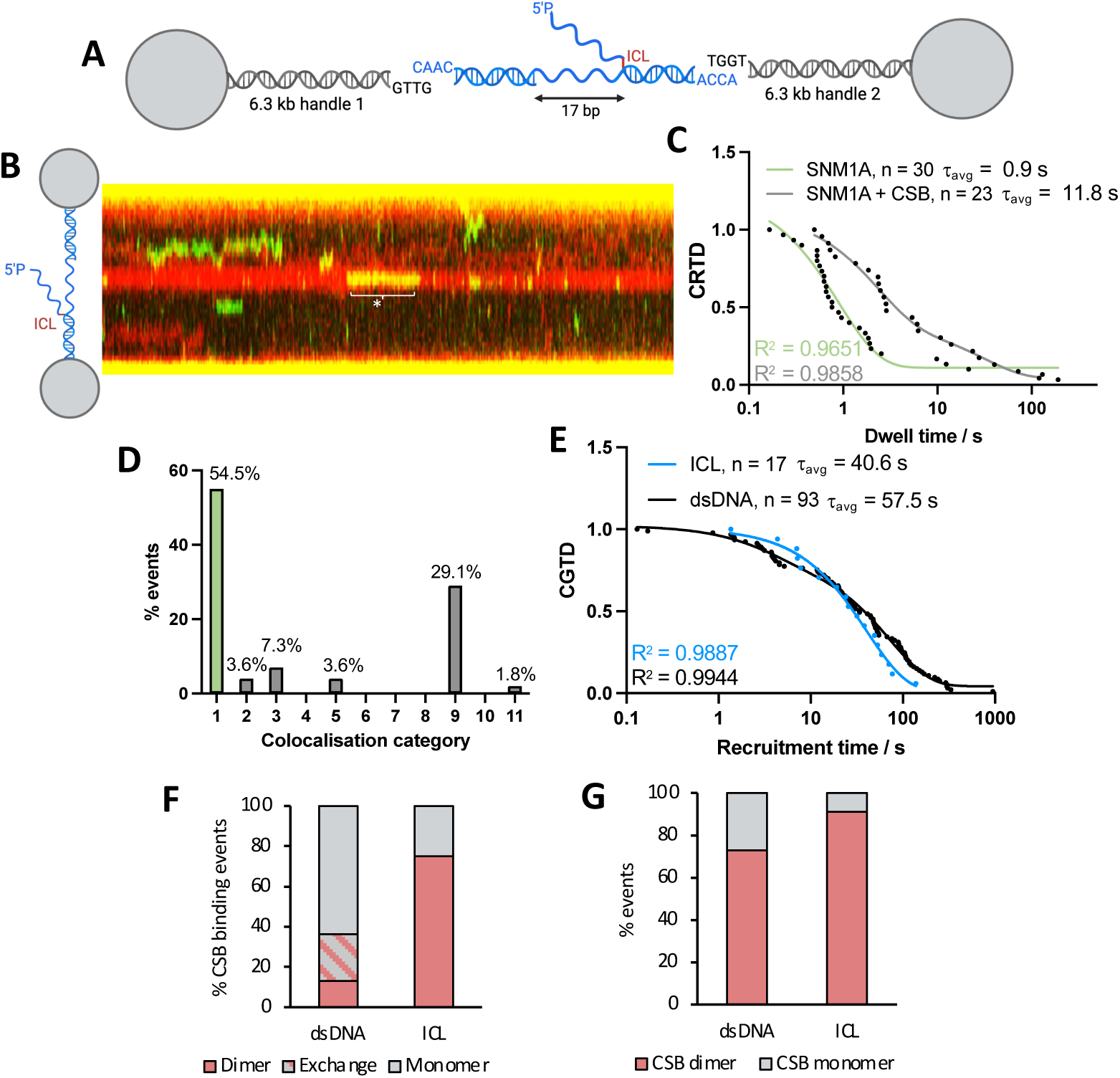
Single-molecule studies of CSB:SNM1A assembly on ICL-containing DNA. **A.** Diagram showing the design of an transcription-coupled ICL repair DNA repair intermediate oligonucleotide that was ligated into two, terminally biotinylated 6.3 kb DNA handles *via* complementary overhangs. **B.** Representative kymograph of CSB (JF-635, red) and SNM1A (JF-552, green) binding to the ICL flap substrate. **C.** CRTD plot of SNM1A dwell times alone (green) or colocalised with CSB in category 9 events (grey) on the ICL. Two-phase binding regimes were determined for colocalised event (with the average given) with both lifetimes given in Table S6. A one-phase binding regime was fit to the SNM1A binding events with the average binding lifetimes shown here. **D.** Bar graph showing the percentage of events represented by different categories of colocalisation for CSB and SNM1A binding alone or together to the centralized ICL. **E.** CGTD plot of recruitment times of SNM1A to CSB-bound DNA on dsDNA (black) or the ICL (blue). A two-phase decay was fit to the recruitment of SNM1A to CSB, with the average given here (both given in Table S6) and a single regime fit to SNM1A recruitment to CSB on ICLs. **F.** Bar graph showing the oligomerization state of all CSB events observed on lambda (ds) DNA or localised at the ICL. Monomer/dimer = CSB event was solely monomeric/dimeric for the entire DNA binding event. Exchange = the CSB oligomerisation state exchanges during DNA binding. **G**. Bar graph showing the oligomerisation state of CSB when co-localised with SNM1A on lambda (ds) DNA or localised at the ICL.

**Figure 7:**
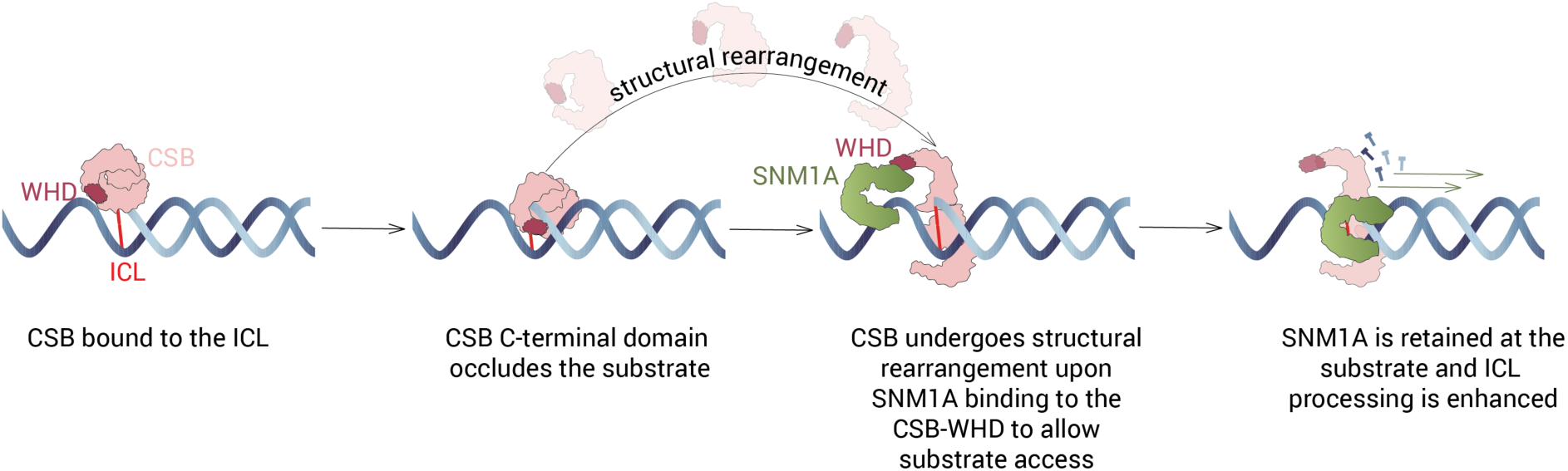
Working model of how CSB enhances the ICL processing activity of SNM1A.

## Discussion

Human SNM1A plays an important role in both replication-dependent and replication-independent ICL repair^46^. In budding yeast, most ICL repair depends upon the ancestral SNM1A homolog, Pso2, and occurs in a manner that is genetically epistatic with, and therefore predicted to operate in the same pathway as, NER factors^47^. By contrast, replication-coupled ICL repair in mammals is controlled through the FA pathway and a number of key NER factors (with the notable exception of XPF-ERCC1) are apparently dispensable within this pathway^48^.

Prior work has identified a physical interaction between SNM1A and CSB, suggesting a potential role for SNM1A in TC-ICL repair^10^, a pathway that has been minimally explored. In support of this concept, CSB is involved in the cellular recruitment of SNM1A to localised, UV-induced psoralen ICL damage ‘stripes’ in a manner dependent upon active transcription. Moreover, in non-dividing neuronal cells, CSB is important for achieving normal levels of ICL unhooking. Together, these observations provide cellular evidence for a coupled role between for SNM1A and CSB in TC-ICL repair, but the nature and molecular details of the interaction between these two proteins and how this interaction influences ICL processing activities of SNM1A has remained unknown.

Our work details the molecular basis of CSB mediated stimulation of SNM1A nuclease activity on ICLs. The results align well with the predicted cellular roles of the CSB:SNM1A interaction^28^. While CSB has been reported to interact with a broad range of DNA repair and transcription factors, influencing their localisation and turnover^14^, an active involvement for CSB in modulating DNA repair enzyme activity has been restricted to several factors involved in BER (NEIL1, NEIL2 and APE1)^18,49,50^ Notably, in no case, are the molecular basis for these CSB-mediated effects understood. Importantly, our single-molecule studies on a single ICL lesion demonstrates that CSB recruits SNM1A to the ICL damage. This is the first visualization of the interaction between CSB and a repair protein interaction in real-time, including on a DNA damage site in a repair-relevant manner. Notably, SNM1A has a 10-fold higher affinity for the damage site when bound by CSB, primarily due to increased binding lifetimes of the SNM1A nuclease (decreased k_off_). This effect is not observed on undamaged, dsDNA, a poor SNM1A substrate. These data support a regulatory role for CSB in DNA damage repair, whereby SNM1A activity is enabled through direct association with CSB.

AF3 modelling of the CSB:SNM1A complex, and subsequent validation by HDX-MS and SPR and immunoprecipitation experiments, provide further insight into how the two proteins interact. In brief, primarily the DNA binding catalytic domain of SNM1A (β-CASP domain) interacts with the WHD of CSB. High affinity binding (equivalent to that observed with full-length CSB) is conferred by the presence of the more extensive C-terminal region of CSB, likely involving the WHD. A higher affinity of SNM1A to DNA-bound CSB versus in-solution CSB, potentially reflects structural rearrangements of the CSB protein that favours SNM1A binding, perhaps by improving accessibility of the WHD. This is also consistent with our nuclease assays which suggested that CSB is already bound to the DNA, modulating, and occluding the substrate, where a specific interaction with SNM1A is required for SNM1A loading onto the substrate. The WHD of CSB has been suggested to mediate other CSB protein-protein interactions, e.g., with RIF1, a downstream effector of 53BP1 that promotes non-homologous end joining in G1^51^. The WHD of CSB is reported to non-covalently interact with monoubiquitin, an interaction for which the biological role is unclear but hypothesised to be required for association of CSB with ubiquitinated repair factors (e.g. CSA or RNAPII)^52,53^. We note here that the affinity between SNM1A and the CSB WHD is over two orders of magnitude greater than the reported interaction between the CSB WHD and monoubiquitin (89 nM vs. 30 μM)^53^, supporting a biologically relevant role for it. WHDs in proteins have also been shown to have functional roles beyond mediating protein-protein interactions, particularly in enabling specific protein-DNA interactions and in aiding strand separation by DNA helicases (such as for RecQ family helicases) ^54^.

We identified and validated a single amino acid variant of CSB, T1447A, that disrupts the interaction with SNM1A and which does not disrupt the interaction with RNAPII. Notably, the interaction between the WHD and SNM1A is necessary, but not sufficient, for stimulation of SNM1A nuclease activity, implying that further communication, either between proteins or with the DNA substrate, are induced by full-length CSB. Indeed, biochemical studies of SNM1A digesting through an ICL showed that while T1447A-CSB does not interact with SNM1A, enhanced SNM1A nuclease activity is still observed in its presence. This suggests that WT CSB can facilitate SNM1A digestion of the ICL, potentially by loading SNM1A onto the DNA substrate, not only via the WHD, but also probably via substrate manipulation^32^. Our DNA-binding studies, indeed, found that the C-terminal domain of CSB is a DNA binding domain, which has not been reported previously. Examination of SNM1A nuclease activity with the C-terminal domain of CSB alone demonstrated that this CSB region partially occludes the DNA substrate, indicating that substrate manipulation and protein-protein coordination is likely necessary after initial CSB:DNA complex formation. For the WHD of CSB, previous co-localisation experiments in cells over-expressing CSB domain constructs demonstrate that the C- and N-terminal CSB domains interact with the core ATPase domain and that disengagement of the WHD from the ATPase domain of CSB is required for efficient NER^55^. Thus, we predict that structural rearrangements of the CSB protein are necessary to unmask the WHD of CSB to orchestrate SNM1A substrate loading and recruitment. Further structural studies will be needed to confirm the above ideas and define the underlying molecular contacts mediating these multiple reactions. Our working model of the basis of CSB enhancement of SNM1A processing is given in Fig 7).

Our work implies a role of CSB dimerisation in DNA repair. While MP experiments show that purified CSB is monomeric in solution, our single-molecule studies suggest that CSB can dimerise on DNA and that this state is strongly favoured on substrates that mimic repair intermediates. This could be attributed to increased protein crowding (as studies were performed in nuclear extracts), and since this would also be the case for our SEC studies where protein concentrations are significantly higher, we do not believe this to be the main cause. We propose that CSB is primarily monomeric in solution but dimerises on DNA upon engagement with preferred structures, as is the case for other transcription factors^56^. In a two colour single molecule experiment, one copy of DNA-bound CSB was shown to recruit another copy. However, on the ICL repair intermediate, dimeric CSB predominated, perhaps reflecting a preference for the in-solution dimer binding to such DNA structures compared with the monomer. Additionally, SNM1A preferentially bound to dimeric, DNA-bound CSB. Roughly 35% of CSB events alone on dsDNA exhibited dimerization, whereas ∼70% of SNM1A events that were colocalised were with the dimeric CSB. This effect was almost complete on ICL repair intermediates, i.e., ∼90% of SNM1A:CSB events were dimeric. This finding contrasts recent cryogenic electron microscopy (cryoEM) structures of CSB in complex with RNA polymerase II and other TC-DNA repair factors in TC repair initiation complexes. In this situation, only one copy of CSB was present ^57,58^. Thus, the apparent descrepancy could reflect an additional, unique biological mechanism of CSB in TC-ICL repair in stimulating SNM1A activity (compared with classical NER where SNM1A is not required), perhaps reflecting a regulatory stage in ICL processing. The cryoEM structures reported revealed only the ATPase domain of CSB, and the complexes studied were crosslinked with glutaraldehyde, providing potential complications that could have resulted in only one copy being present or observed due to filtering during particle processing.

It is interesting to note that SNM1A, a potent and potentially damaging 5′-3′ exonuclease, has a relatively low affinity for all DNA substrates examined to date, but has high affinity for a number of DNA binding ligands (e.g. CSB) that can direct its activity to relevant repair intermediates. In replication-coupled ICL repair, binding of monoubiquitinated PCNA is required through the interaction of the SNM1A PIP box and an N-terminal UBZ (ubiquitin binding domain)^64^. Moreover, for the recently identified role of SNM1A in the repair of complex DNA double-strand breaks, a PAR-binding zinc finger (PBZ) protein appears to act in concert with the PIP box and UBZ motif to control recruitment of SNM1A in anticipation of its repair role^65^. How SNM1A is subsequently released from sites of repair and kept in a functionally dormant state, where it is not able to access non-pathological, free DNA 5′ termini, remains unknown and is a topic of substantial interest for future investigation.

While the interaction between SNM1A and the CSB C-terminus is of high affinity and is required for efficient localisation of SNM1A to sites of TC-ICL repair, it remains uncertain what other factors are involved in the process of TC-ICL repair^20^. By analogy with TC-NER, it will be important to assess the involvement of other core NER factors and accessory TC-NER factors (e.g., CSA, UVSSA, ELOF1, STK19, etc.) reviewed in^14,66,67^. The recent evidence for the involvement of UVSSA, CSA and TFIIH components in TC-ICL repair is also noteworthy^68^. Additionally, canonical TC-NER involves numerous post-translational modifications (PTMs) and other interactions, including ubiquitination of CSB by CSA ^69^ and RNA Pol II ubiquitination^67,70,71^. As research on the molecular details of TC-ICL repair is in its infancy, building a comprehensive picture of the cellular and molecular requirements is needed, including the spatial and temporal relationship with other transcription and repair factors and the impact of PTMs. It is worth emphasizing that the barrier to RNA Pol II progression presented by an ICL is structurally distinct from lesions more widely studied (e.g. UV-induced damage) that are repaired by TC-NER. ICLs could arrest RNA Pol II progress at a greater distance from the lesion due to their ability to prevent strand separation during transcription, and therefore assumptions of similarity with classical TC-NER should be made with caution.

The potential physiological importance of TC-ICL repair is underscored by recent studies examining the interplay between aldehyde detoxification and transcription-associated DNA repair^72^. In these studies, the cellular accumulation of formaldehyde (genetically induced, *via ADH5* knockout) produced endogenous lesions that blocked transcription (noting that formaldehyde can produce a wide range of DNA lesions, including both ICLs and DNA protein crosslinks^73^). Subsequent elimination of CSB in these *ADH5^-/-^* mice induced features reminiscent of Cockayne syndrome. It is of interest to determine whether, at a physiological level, the key TC-ICL repair unhooking step we have begun to define here, catalysed by CSB and SNM1A, plays a key role in removing the endogenous DNA damage that contributes to premature aging features and neurodegeneration seen in Cockayne syndrome-affected individuals and the population as a whole.

DDR nucleases have long been targeted for cancer treatment and some advances have been made; however, sufficiently safe, selective, and potent active site targeting inhibitors suitable for *in vivo* target validation have yet to be identified. Our results suggest that targeting protein-protein interactions that are vital for specific nuclease activities, such as is the case with CSB mediated stimulation of SNM1A’s nuclease activity, may be a productive alternative to active site targeting^74,75^. Since many nucleases have multiple context-dependent functions, such an approach may be particularly useful for achieving selectivity with respect to disruption of specific aspects of nuclease mediated biological processes, e.g., only the TC-ICL repair role of SNM1A.

## Acknowledgements

We thank Dr. David Staunton in the Molecular Biophysics Suite, Department of Biochemistry, University of Oxford for assistance with circular dichroism spectroscopy, Dr. Rod Chalk from the Centre for Medicines Discovery, University of Oxford for mass spectrometry analysis and Dr. Gemma Harris at Harwell Research Campus for performing SEC-MALS experiments. This work was supported by Cancer Research UK Discovery Programme awards CRUK/A24759 and DRCRPG-Nov23/100005 to PJM and CJS, an NIH-OxCam student award to WJN, a Wellcome Trust award 218514/Z/19/Z to RvW and the NIH R35ES031638 (BVH), F32ES034982 (MAS) and S10OD032158 (C-Trap). Molecular graphics and analyses were performed with UCSF ChimeraX, developed by the Resource for Biocomputing, Visualization, and Informatics at the University of California, San Francisco, with support from National Institutes of Health R01-GM129325 and the Office of Cyber Infrastructure and Computational Biology, National Institute of Allergy and Infectious Diseases. We also thank Dr. Luke Lavis (Janelia Research Campus) for the HaloTag ligands used. Figures were generated in BioRender.

## Data availability

Python scripts and methods used for single-molecule analyses are detailed in Rakowski *et al*.^76^. Raw data is available upon request.

## Methods

### Plasmids for recombinant protein expression

Full-length DNA encoding for CSB (*ERCC6*) was acquired as a synthetic gene from Genscript in the pFastBac1 vector, codon optimized for *Spodoptera frugiperda* (*Sf*9) expression with N-terminal His_10_ and ZBasic tags with a TEV protease cleavage site and a C-terminal Avi tag. For preparation of biotinylated protein, the full-length CSB gene (including tags) was subcloned from the pFastBac1 vector by InFusion cloning into a pFastBac Dual vector, with CSB expressed under the polyhedrin promoter and BirA (with C-terminal mCherry tag) co-expressed under the p10 promoter. Plasmids encoding CSB lacking the N- or C-terminal domains (ΔN-CSB, a.a. 402-1493 and ΔC-CSB a.a. 1-1014) were cloned using InFusion from the full-length CSB vector. Production of N-Avi-SNM1A_698-1040_ was performed using the same pFastBac Dual vector (co-expressing BirA-mCherry), with the SNM1A sequence codon optimized for baculovirus expression.

The pNIC28-Bsa4 vector (with N-terminal His_6_ followed by TEV cleavage site) was used for production of all C-terminal and N-terminal CSB domains, with the CSB sequence codon optimized for expression in *E. coli*. For expression of the ATPase domain of CSB (residues 402-1040), codon optimization for baculovirus expression and ligation independent cloning was used to transfer to the pFB-LIC-Bse expression vector, giving a fusion protein with N-terminal His_6_ followed by a cleavage site. Single-point mutations (SNM1A-E864A, CSB-T1447A and CSB-E647Q, CSB-F796A) were generated from the wild-type plasmids, using InFusion cloning.

For single-molecule studies, codon-optimized mammalian full-length CSB and C-terminal SNM1A sequences (a.a. 698-1040) were subcloned into vectors with N-terminal HaloTags under a CMV promoter (Promega: pFN28A and Addgene: 58241).

All plasmids generated were verified by Nanopore sequencing (Cambridge Biosciences).

### Expression, and purification of recombinant proteins

For bacmid preparation, full-length and ATPase domain (a.a. 402-1040) CSB plasmids were transformed into DH10Bac *E. coli* cells (Life Technologies). Bacmid DNA was isolated and used to transfect *Sf*9 insect cells (Gibco^TM^, cultured in Sf-900 II SFM). Viral amplifications were performed following the manufacturer’s instructions. For expression, *Sf*9 cells were prepared at 2-3 x 10^6^ cells/mL, 7 mL/L culture of P2 virus was added and incubated at 27 °C for 72 h at 180 rpm. For expression of the biotinylated full-length CSB (in pFastBac Dual vector co-expressing BirA-mCherry), 50 μM D-biotin was added (1 in 1000 of 50 mM stock prepared in 0.1 M NaOH). The cells were harvested by centrifugation (1,500 x *g*, 10 min, 4 °C) and stored at -80 °C until further processing. For the CSB ATPase domain, purification was performed as per the C- and N-terminal fragments (expressed in *E. coli* and described later). For full-length CSB, lysis buffer (80 mL per L of initial culture; 50 mM HEPES pH 8, 1 M NaCl, 20 mM imidazole, 1 mM tris(2-carboxyethyl)phosphine (TCEP), 10% *v/v* glycerol + 1 Roche cOmplete EDTA-free protease inhibitor tablet) was used to resuspend the pellet on ice. Cells were lysed by sonication (10 A, 6 x 30 s ON, 60 s OFF) then centrifuged (74,000 x *g*, 60 min, 4 °C). The supernatant was filtered (0.45 μm), then loaded onto Ni-NTA agarose beads (1 mL beads per L of expression culture pre-equilibrated with lysis buffer). The beads were washed with lysis buffer (20 CV), lysis buffer + 750 mM NaCl + 40 mM imidazole (20 CV) and the protein was eluted with 5 CV of lysis buffer + 500 mM NaCl + 300 mM imidazole with 5% *v/v* glycerol (vs. 10% *v/v* glycerol previously). Protein-containing fractions (the 300 mM imidazole elution) were pooled, desalted into gel filtration buffer (50 mM HEPES pH 8, 300 mM NaCl, 1 mM TCEP) then loaded onto a Superdex S200 GL 10/300 column (pre-equilibrated with gel filtration buffer). Protein-containing fractions were pooled (protein elutes 9.5-13.5 mL) and diluted 2-fold with a salt-free buffer (50 mM Tris pH 8, 1 mM TCEP). The protein was loaded onto a 1 mL HiTrap SP FF column; an elution gradient was performed from 150 mM NaCl to 1.5 M NaCl over 8 column volumes at 1 mL/min. Protein-containing fractions were pooled, concentrated, aliquoted and stored at -80 °C.

N-Avi-SNM1A_698-1040_ (WT and E864A) bacmid DNA was prepared as per CSB expressions and was produced in *Sf9* insect cells prepared at 2-3 x 10^6^ cells/mL then 7 mL/L of culture of P2 virus and 500 μL/L of culture of D-biotin (50 mM in 0.1 M NaOH) were added. Cells were incubated at 27 °C for 72 h at 180 rpm. The cells were harvested by centrifugation (1,500 x *g*, 10 min, 4 °C), washed with 1 x PBS (4 °C), then stored at -80 °C until further processing. The pellet was resuspended in lysis buffer (80 mL per L of initial culture: 20 mM HEPES pH 7.5, 500 mM NaCl, 5% v/v glycerol, 0.5 mM TCEP + 1 Roche cOmplete EDTA-free protease inhibitor tablet) on ice. Cells were lysed by sonication (10 A, 6 x 30 s ON, 60 s OFF) then centrifuged (74,000 x *g*, 60 min, 4 °C). The supernatant was incubated with Streptavidin mutein matrix beads (1 mL, Roche) for 1 h, 4 °C, with gentle rolling. The unbound protein was removed by centrifugation (3,000 x g, 5 min, 4 °C) and the beads transferred to a gravity column. The beads were washed with lysis buffer (50 mL) and then the protein eluted in 5 mL lysis buffer + 5 mM D-biotin. If tags were not to be removed, the elution was concentrated and injected onto a Superdex S75 GL 10/300 column (in 20 mM HEPES pH 7.5, 300 mM NaCl, 0.5 mM TCEP). Protein containing fractions were pooled, concentrated, and stored at -80 °C.

If the biotinylated Avi tag was removed, the protein was incubated with TEV protease overnight with dialysis into gel filtration buffer and gel filtration was performed as with the tagged protein.Protein containing fractions were pooled, concentrated, and stored at -80 °C. Expression and purification of N-His-SNM1A_698-1040_ has already been reported ^77^.

Plasmids for CSB_1187-1493_, CSB_1308-1493_, CSB_1424-1493_, CSB_1187-1401_, and CSB_1187-1308_ were expressed in BL21(DE3) pLysS cells. A single colony from a fresh transformant was used to inoculate LB media (+ 50 μg/mL kanamycin sulfate final). This culture was incubated overnight at 37 °C, 220 rpm then added at 2% *v/v* final to 1 L of LB media (+ 50 μg/mL kanamycin sulfate final). The culture was incubated at 37 °C, 220 rpm until OD_600_ = 10-20 then cooled to 18 °C and induced by addition IPTG (to 0.5 mM final). The culture was incubated at 18 °C for 16-18 h. Cells were harvested by centrifugation (5,000 x *g*, 15 min, 4 °C), then stored at –20 °C until purification. For protein purification (for all C-terminal, N-terminal and ATPase domain fragments), cells were resuspended in lysis buffer (40 mL per L of expression culture; 50 mM HEPES pH 8, 500 mM NaCl, 5% glycerol, 0.5 mM TCEP + 1 cOmplete EDTA-free protease inhibitor tablet) and sonicated (16 A, 3 x 1 min ON, 1 min OFF) on ice then centrifuged (50,000 x *g*, 4 °C, 30 min). The supernatant was incubated with 2 mL Ni-NTA agarose beads for 30 min, 4 °C, with gentle rolling. The unbound protein was removed by centrifugation (3,000 x *g*, 5 min, 4 °C) and the beads transferred to a gravity column. The beads were washed with lysis buffer (30 mL), lysis buffer + 40 mM imidazole (30 mL) then the protein eluted with lysis buffer + 300 mM imidazole (5 mL). The protein was desalted into gel filtration buffer (50 mM HEPES pH 8, 300 mM NaCl, 0.5 mM TCEP) using Zeba Spin 7K MWCO desalting columns and injected onto a Superdex S75 GL 10/300 column (pre-equilibrated with gel filtration buffer). Protein-containing fractions were analysed by SDS-PAGE and those containing the desired protein were pooled, concentrated, flash-frozen and stored at -80 °C.

For preparation of the CSB:SNM1A complexes, comprising of CSB_1424-1493_, CSB_1187-1493_ or full-length CSB, and N-Avi-Biotin-SNM1A_698-1040_, the two proteins were individually expressed and incubated with affinity chromatography columns as previously described (Ni-NTA agarose and streptavidin mutein matrix, respectively with all buffers supplemented with 1 mM Mg^2+^). The SNM1A-bound streptavidin mutein matrix beads (1 mL per 500 mL expression culture) were washed with SNM1A lysis buffer. CSB bound to Ni-NTA was washed, eluted, and desalted (as previously described for the protein alone) and incubated with SNM1A bound to streptavidin mutein matrix beads for 30 min, 4 °C. Unbound protein was removed from the beads by centrifugation (3,000 x *g*, 4 °C, 5 min) and the beads were washed with gel filtration buffer (20 CV, 20 mM HEPES pH 8, 500 mM NaCl, 0.5 mM TCEP, 1 mM Mg^2+^) in a gravity column. The protein was eluted with 5 CV gel filtration buffer + 5 mM D-biotin. The protein was concentrated and injected onto a Superdex S200 GL 10/300 column. Protein-containing fractions were analysed by SDS-PAGE and those containing the desired protein were pooled, concentrated, flash-frozen and stored at -80 °C.

### Oligonucleotide preparations

High-performance liquid chromatography (HPLC)-purified oligonucleotides used for biochemical assays were purchased from Eurofins Genomics. Stocks were prepared at 100 μM in nuclease-free water and stored at -20 °C. Oligo sequences and annealing partners to give different secondary structures are given in Table S3 and Fig. S8.

### Preparation of ICL-containing oligonucleotides

The ‘0-nucleotide substrate’ and C-Trap substrate were synthesized by phosphoramidate chemistry with a triazole crosslink formed by a Cu(I)-catalysed click chemistry reaction between azide and alkyne groups incorporated on opposing bases in the duplex. Synthesis was performed as reported^78^. Briefly, both strands of the respective DNA substrates were synthesised independently using an Applied Biosystems 394 DNA/RNA Synthesizer. Solid-phase synthesis was performed using a standard 1 µmol phosphoramidite cycle of acid-catalysed detritylation, coupling, capping, and iodine oxidation. The coupling time for normal nucleoside phosphoramidite monomers (A, T, G, C) was 1 minute. For the azide-containing nucleotide, 5’-(4,4’-dimethoxytrityl)-N6-benzoyl-N8-[6-(trifluoroacetylamino)-hex-1-yl]-8-amino-2’-deoxyadenosine,3’-[(2-cyanoethyl)-(N,N-diisopropyl)]-phosphoramidite (‘amino-modified C6 dA,’ Glen Research) was incorporated (coupling time: 14 minutes). For the alkyne-containing nucleotide, 5’-dimethoxytrityl-5-(octa-1,7-diynyl)-2’-deoxyuridine,3’-[(2-cyanoethyl)-(N,N-diisopropyl)]-phosphoramidite (‘C8-Alkyne-dT-CE Phosphoramidite’, Glen Research) was incorporated (coupling time: 14 minutes). Following synthesis completion, oligonucleotides were cleaved from the solid support and deprotected by exposure to concentrated aqueous ammonia solution (1 h, rt), then heated within a sealed tube (5 h, 55°C). Oligonucleotides were purified using HPLC with triethyl ammonium bicarbonate buffer (TEAB), as reported ^79^.

To convert the amino-modified C6 dA amino group into an azide, the oligonucleotide (200 nmol) was prepared with 6-azidohexanoic acid NHS ester (1 mg, 3.9 mmol) in 1:1 dimethyl sulfoxide: 0.5 M Na_2_CO_3_/NaHCO_3_, pH 8.75 (160 μL) for 4 h at rt. The azide-labelled oligonucleotide was then purified by reversed-phase HPLC, as previously reported ^79^. Under an inert atmosphere, the alkyne and azide strands were prepared in 0.2 M NaCl solution (110 μL, 50 μM) and annealed by heating to 95 °C then allowing to cool to room temperature (2-3 h). Under an inert atmosphere, CuSO_4_.5H_2_O (0.03 mg, 0.12 μmol from an 8 mM stock in H_2_O), Tris(3-hydroxypropyltriazolylmethyl)amine ligand (0.5 mg, 1.15 μmol) and sodium ascorbate (0.2 mg, 1.0 μmol from a 500 mM stock in H_2_O) were prepared in water (15 μL). This was added to the annealed DNA and incubated for 2 h at rt. The completed click reaction was then desalted using a NAP-25 DNA purification column (Cytiva) and lyophilised.

Desalted click reactions were purified by denaturing PAGE using an 8% 19:1 acrylamide:bis-acrylamide and 7 M urea gel. Successfully crosslinked DNA substrate migrated more slowly compared to non-crosslinked strands (Fig. S10). Crosslinked substrates were cut out from the gel and recovered by the crush and soak method. The gel pieces were crushed and soaked in 30 mL of water for 18 hours at 37 °C while shaking. Gel particles were filtered out and the resulting solution was desalted using a NAP-25 column, lyophilised and resuspended in water. The correct identity of the final crosslinked products were confirmed by quadrupole time-of-flight (Q-TOF) mass spectrometry (Fig. S10).

### Generation of 3**′** radiolabelled oligonucleotides

All substrates, except the ‘0-nucleotide’ (herein referred to as ‘0-nucleotide’) ICL-containing substrate, were 3′-labelled using terminal deoxynucleotide transferase (TdT) (NEB), then annealed with the complementary oligonucleotide. For labelling, oligonucleotide (10 pmol), TdT (20 U) and α-^32^P-dATP (3.3 pmol) were prepared in 1 x TdT buffer (NEB) and incubated at 37 °C for 1 h. The reaction mixture was then passed through a P6 Micro Bio-Spin chromatography column (Bio-Rad, 1,000 x *g*, 4 min, rt), to give the labelled oligonucleotide at ∼500 nM. To generate duplex-containing substrates, the annealing reactions were performed with radiolabelled substrate: unlabelled substrate(s) 1:1.5 molar ratio by heating to 100 °C for 5 min in annealing buffer (10 mM Tris pH 7.5, 100 mM NaCl, 0.1 mM EDTA) then allowing to cool to room temperature for 3-18 h. Stocks were prepared at 100 nM final based on the radiolabelled oligonucleotide.

For labelling of the ‘0-nucleotide’ ICL substrate, the oligonucleotide (10 pmol) was first annealed by boiling to 100 °C for 5 min, then allowed to cool to rt in annealing buffer. The oligonucleotide (10 pmol) was incubated with Klenow (exo-) (12.5 U, NEB) and α-^32^P-dATP (3.3 pmol) in 1 x NEB buffer 2 at 37 °C for 45 min. dGTP (1 mM) and dATP (1 mM) were added to 0.1 mM final concentrations and the mixture incubated for 30 min at 37 °C. The reaction was then passed through a P6 Micro Bio-Spin chromatography column (Bio-Rad, 1,000 x g, 4 min, rt), annealed in 1 x annealing buffer and prepared at 100 nM oligonucleotide concentration.

### Nuclease assays

Reactions were prepared in 20 mM HEPES KOH pH 7.5, 50 mM KCl, 4 mM MgCl_2_, 0.05%*_v/v_* Triton X-100, 0.5 mM TCEP, 5% *v/v* glycerol. Protein was added first, followed by 10 nM oligonucleotide (all concentrations given are final) on a 10 μL scale. Reactions were incubated at 37 °C for the time indicated, then reactions quenched by addition of stop solution (10 μL; 95% formamide, 10 mM EDTA, 0.25%*_v/v_* xylene cyanol, 0.25%*_v/v_* bromophenol blue) then incubated at 100 °C for 5 min. Samples were then immediately incubated at 4 °C for 5 min, then centrifuged (1 min, 13,000 x g, 4 °C). Analysis was performed by 20% denaturing PAGE (50% *v/v* 40% 19:1 acrylamide:bis-acrylamide, 7 M urea, 1 x tris-borate EDTA, 0.12% *w/v* ammonium persulfate, 0.12% *v/v* tetramethylethylenediamine). The gel was preheated (700 V, 1 h) then electrophoresis performed at 700 V, 1 h. Gels were fixed for 1 h in 50% *v/v* methanol, 10%*_v/v_* acetic acid in water, dried at 80 °C for 2 h under vacuum, then exposed to a phosphor imager screen (Kodak). Screens were scanned with a GE Typhoon FLA 9500. Quantification was performed in ImageJ, measuring the amount of intact substrate, compared to that which had been digested. Assays were quantified based upon triplicate reactions, with the end of each error bar representing each data point.

### Electrophoretic mobility shift assays (EMSAs)

Reactions were prepared on ice in 20 mM HEPES KOH pH 7.5, 50 mM KCl, 0.05% *v/v* Triton X-100, 0.5 mM TCEP, 5%*_v/v_* glycerol, 1 nM oligonucleotide and the indicated concentration of protein on a 10 μL scale. All concentrations given are final. The metal ion or EDTA concentration used, if applicable, is given in each relevant figure. For the binding of a single protein to DNA, protein was added last to the reaction whereas to study potential for co-operative binding of proteins to DNA, DNA was added last. Reactions were incubated at 4 °C for 10 min, then immediately loaded onto a 0.5% agarose gel (in 0.5x TBE, pre-cooled to 4 °C). The gel was run at 100 V for 90 min at 4 °C then fixed (30 min - 1 h in in 50% methanol, 10% acetic acid) and dried under vacuum (50 °C, 1 h) on three pieces of Whatman filter paper. The gel was then exposed to a phosphor imager screen (Kodak) and imaged using a GE Typhoon FLA 9500.

### Surface plasmon resonance (SPR) assays

Protein-protein interactions were measured using a Biacore S200 machine using a Series S SA sensor surface. SNM1A was immobilised via a biotinylated N-terminal Avi-tag at an immobilisation level of ∼4000 RU. Binding analysis using CSB analytes was performed in a buffer containing 20 mM HEPES pH 7.5, 150 mM NaCl, 0.5 mM TCEP. Single cycle analysis was performed using either 5 or 6 two-fold dilutions starting from 500 nM or 1 μM. CSB protein was injected for 90s with the final dissociation phase of at least 900 s. Full-length CSB was immobilised *via* a biotinylated C-terminal Avi tag in 20 mM HEPES pH 7.5, 500 mM NaCl, 0.5 mM TCEP, 0.01% TWEEN-100. SNM1A was injected for 90 s with the final dissociation phase of at least 900 s.

For multi cycle analysis SNM1A was immobilised *via* a biotinylated N-terminal Avi-tag at an immobilization level of approximately 4000 RU. Two-fold serial dilutions starting from 2 μM were injected for 300 seconds with a dissociation phase of 1200 seconds. Regeneration was performed between each injection using the mobile phase buffer containing 1 M NaCl.

Data were fitted using Biacore S200 evaluation software. Single cycle curves were fitted using a 1:1 Langmuir binding model with accommodation for drift as a single parameter when necessary (based upon an estimation of the curve before the first injection). Fitting parameters are give in Supplementary Tables 4 and 5.

### Circular Dichroism (CD) spectroscopy

Samples were prepared at 0.1 mg/mL (determined by Nanodrop spectrophotometer) in 10 mM sodium phosphate pH 7, 20 mM NaCl in a 0.1 cm quartz cuvette. Measurements were performed using a JASCO-815 spectropolarimeter with a Peltier temperature control unit. The CD signal (in mdeg) was measured at wavelength 195-250 nm at 20 °C.

### AlphaFold3

AlphaFold3 used the opensource online server with standard presets ^80^. USCF ChimeraX1.15 was used to visualize the results and generate figures^81^. PICKLUSTER (via USCF Chimera X1.5 plug-in) was used to analyse AlphaFold3 model interfaces ^82^.

### Hydrogen-deuterium exchange mass spectrometry

*Sample Preparation:* To investigate the SNM1A interaction interface with CSB, Hydrogen Deuterium eXchange Mass Spectrometry (HDX-MS) experiments were performed. Initial CSB stocks were at 9 μM in 50 mM HEPES, 300 mM NaCl, 0.5 mM TCEP and in the absence of glycerol. ??OK?? SNM1A was at 55 μM in 50 mM HEPES, 300 mM NaCl, 0.5 mM TCEP, 5% glycerol buffer. CSB was incubated in the presence (holo-state) and absence (apo-state) of SNMIA for 60 min at room temperature to facilitate full complex formation, with a CSB:SNMIA molar ratio of 1:2.5. Samples were diluted to achieve a final concentration of 20 pmol on column for CSB.

*Data Acquisition:* HDX-MS was performed as reported^83^. Briefly, a labeling buffer mimicking that of the protein stock conditions were prepared in deuterium oxide D_2_O (99+ %D, Cambridge Isotope Laboratories, Tewksbury, MA), with the pH adjusted to pD 7.50 (pD = pH+0.4). The quenching buffer comprised of 0.8**%**_v/v_ aqueous formic acid (FA), pH 1.08 in H_2_O. Following initial complex formation and equilibration, samples were diluted in deuterium labelling buffer in a 1:10 ratio to achieve a final excess D_2_O concentration of 90%. Labelling time points of 0.5, 10, and 60 min were sampled at 20°C, in addition to non-deuterated controls in H_2_O buffer. All analyses were acquired at least in triplicate. Following completion of each labeling time, samples were quenched by adding quench buffer (1:1 ratio), to acheive a final pH of 2.51. Samples were digested using a column (2.1 x 3.0 mm; NovaBioAssays, MA) containing a combination of pepsin and proteaseXIII acidic proteases at 8°C for 3 min. Peptides were then trapped on a 1.0 mm x 5.0 mm, 5.0 µm trap cartridge (Thermo Scientific™ Acclaim PepMap100) for desalting and at a flow rate of 150 µL/min. Peptides were separated by increasing hydrophobicity on a Thermo Scientific™ Hypersil Gold™ column (50 x 1 mm, 1.9 um, C18) by a linear gradient of 5% to 40% Buffer B (A: water, 0.1% FA; B: ACN, 0.1% FA) and a flow rate of 100 µL/min. Following each sample injection, a protease wash (of 2M guanidine, 0.8% FA, pH 2.3 in H_2_O was performed to minimise peptide carry-over. To limit back-exchange, the LC system was maintained at 1.5°C. Labeling, quenching, and online digestion steps were facilitated by use of an automated HDX robot from Trajan Scientific and Medical. The acquisition and management of samples were guided by Chronos (version 5.4.1). All labelled samples were acquired in MS1 mode on a Thermo Scientific™ Orbitrap Exploris™ 480 Hybrid™ mass spectrometer.

*Data Analysis:* Prior to the labelling experiment, an unspecific digested database of non-deuterated CSB and SNMIA peptides was created in BioPharma Finder (version 5.2) through a data-dependent and targeted HCD-MS2 acquisition regime. Labeling data were processed and manually curated using HDExaminer version 3.4.2 (Trajan Scientific and Medical). The charge state with the highest quality spectra for all replicates for each peptide across all HDX-MS labeling times was used in the final analysis. Residual plots, Volcano plots, and heat maps were generated to compare both the apo- and holo-states of CSB. Significant differences observed at each residue of the protein was used to map HDX-MS consensus effects (based on overlapping peptides) onto the AlphaFold model of CSB. All MS raw files were uploaded to ProteomeXchange (https://www.proteomexchange.org/) and can be found under the unique identifier PDXXXXXX.

### Mass photometry

The protocol for MP measurement has been described in detail ^84^. Briefly, measurements were recorded on a TwoMP (Refeyn Ltd, Oxford) in silicon gaskets (GBL103250, Grace Bio-Labs) attached to precleaned microscope coverslips (24 × 50 mm, Menzel Gläser, VWR 630-2603), functionalised with (3-aminopropyl)triethoxysilane (APTES, Sigma Aldrich, 440140). Coverslips were cleaned by consecutive 5 min bath sonications in water (Milli-Q® water, 18.2 MΩ·cm), 50% isopropanol in water and water again, and then dried with nitrogen gas. Coverslips were subsequently exposed to oxygen plasma under 0.5 mbar of oxygen at 40% power for three minutes (Zepto plasma cleaner, Diener Electronic). Directly after, the coverslips were incubated in 2% APTES in acetone for three minutes while stirring (Sigma Aldrich, 34850-M). The coverslips were subsequently washed in acetone and placed in a preheated oven at 110 °C for two hours, followed by two times bath sonication in 50% isopropanol in water and then in water. The coverslips were then dried under a flow of nitrogen gas. Buffer was added to the gasket first to adjust the focus position and optimise the contrast at the glass-solution interface. The protein sample was then added to the gasket containing the buffer and a 60 second movie was recorded using the regular field of view of (10.9 x 4.3 μm^2^) and frame rate of 238 Hz (AcquireMP V2023 R1.1, Refeyn Ltd.). Movie processing and event detection was done in DiscoverMP software using a moving window of 5 frames to generate the ratiometric movie (V2024 R1 Refeyn, Ltd.). The measured contrasts were converted to mass using a contrast to mass protein calibrant as described previously ^85^.

All MP measurements and incubations were performed in 50 mM HEPEs pH 7.5, 300 mM NaCl, 0.5 mM TCEP. To measure the oligomeric state of CSB, the protein stock solution was rapidly diluted from 2 μM (determined using UV-VIS) to to 20 nM in the gasket for measurement. To calculate the affinity between CSB and SNM1A, a mixture of 200 nM of both proteins was incubated at room temperature for 5 minutes and rapidly 10-fold diluted into the gasket. To determine the affinity of the CSB:SNM1A interaction, we used the fact that the relative abundances are representative of the partial solution concentration and determined the relative abundances by fitting the peaks in the histogram with a Gaussian function. The dissociation constant was calculated by:

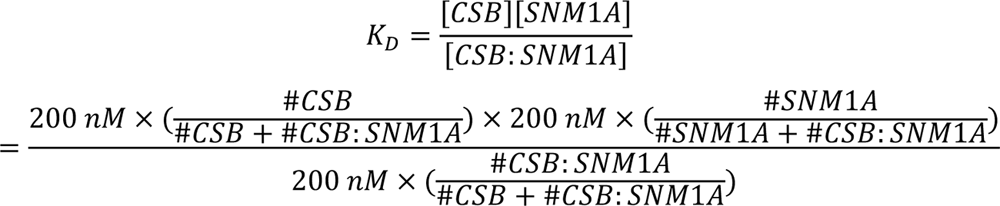

### CSB nuclear extract preparation for single-molecule studies

Expi293F cells were cultured in a 5% oxygen atmosphere using the ??OK?? Expi293^TM^ expression medium. Cells were seeded at 2.4 x 10^6^ cells/mL, transfected with 1 μg plasmid per 1 mL of cells. Transfections were performed on 20 mL of Expi293F cells. Initially, 20 μg of plasmid was diluted in 1 mL OptiMEM, and 60 μL polyethyleneimine (PEI, linear MW = 25 kDa, at 1 mg/mL) prepared in 1 mL OptiMEM. The two solutions were incubated for 5 min at rt, then the PEI solution mixed gently with the DNA mixture and incubated for 30 min at rt. The resulting solution was added to 20 mL Expi293F cells and cultured for 18 h. Valproic acid (3.5 mM final) and non-essential amino acids (ThermoFisher: 11140050, added to 1 X final) were then added. 65 h after initial transfection, JaneliaFluor dye was added to 100 nM final and cultured for 30 min. Cells were harvested by centrifugation (1,000 x g, 15 min, 4 °C) and the supernatant discarded. Nuclear extracts were prepared using 1 mL of cells post-transfection. Cells were washed with 1 x PBS (3 x 1 mL PBS, with centrifugation 1,000 x g, 5 min, 4 °C between each wash) and nuclear extracts prepared using an Abcam nuclear extraction kit (ab113474) following the manufacturer’s protocol. Protein concentrations were determined using the Bradford assay and western blots were used to quantify the amount of HaloTagged protein target vs. unlabelled protein (Fig S21). SDS-PAGE gels were run and imaged based on the JF dye used to determine the percentage of labelled protein compared with free dye (Fig S21). Single-molecule experiments were performed using CSB produced on two separate days using either the Promega or Addgene vector (as desrcibed above). Western blot analysis was performed to investigate whether RBP1 was present in the nuclear extract (Fig S22).

### SNM1A preparation for single-molecule studies

For preparation of partially-purified SNM1A_698-1040_ for single-molecule studies, transfection and labelling was performed in the same manner as for the CSB nuclear extract preparation. The cells (15 mL expression culture) were harvested by centrifugation (1,000 x g, 15 min, 4 °C) and the supernatant was discarded. The pellet was washed with 1 x PBS (5 mL) and resuspended in 10 mL xTractor (Takara) supplemented with 20 mM imidazole (final), 500 mM NaCl (final), DNase I and one cOmplete mini EDTA-free protease inhibitor cocktail tablet, then incubated for 20 min on ice. Cell debris was pelleted by centrifugation (10,000 x g, 20 min, 4 °C) and the supernatant incubated with 100 μL Ni-NTA agarose beads (Qiagen) for 1 h at 4 °C. The beads were washed (1 x 3 mL buffer: 50 mM HEPES pH 7.5, 500 mM NaCl, 20 mM imidazole, 0.5 mM TCEP, 10% glycerol) followed by washing with 1 x 500 μL buffer + 40 mM imidazole and elution with 1 x 100 μL buffer + 300 mM imidazole. The eluted protein was aliquoted and frozen at -80 °C until use. Protein concentrations were determined using a Bradford assay and western blots used to quantify the amount of HaloTagged protein target vs. unlabelled (Fig S21). SDS-PAGE gels were run and imaged based on the JF dye used to determine the percentage of labelled protein compared with free dye (Fig S21). Single-molecule experiments were performed using SNM1A expressed on two separate days using the Addgene vector, giving an N-terminal His-Halo tag (detailed above).

### DNA substrate preparation for single-molecule studies

Biotinylated lambda DNA was prepared as described by Schaich *et al*. 2023 ^42^. To incorporate the ICL-containing substrate into the C-Trap, the LUMICKS’ DNA tethering kit (SKU: 000026-1) prepared adapting the protocol of Schnable *et al.* 2024^86^. The oligonucleotide sequences and resulting structure used are given in Figure S4 and S10. Oligonucleotides were purchased from IDT and the two oligonucleotides containing the alkyne- and azide-containing modifications were cross-linked via ‘click’ chemistry (to form the ICL) and purified, as described in prior methods. The resulting cross-linked oligonucleotide was then annealed with a ‘top’ strand oligonucleotide to give the oligonucleotide at 1 μM final and to provide complementary handles to the LUMICKS’ tethering kit arms. The annealed oligos were then diluted to 25 nM and ligated into the Lumicks DNA tethering kit overnight at room temperature. Ligation success was measured by generating DNA tethers on the LUMICKS C-trap and comparing force-distance curves to the extensible worm-like chain model for 12 kb DNA products (which can only form if both 6 kb handles ligate onto the middle insert). After preparation, the substrate was stored at 4 °C and diluted freshly by a factor of 1/300 in PBS before each day of imaging.

### Single-molecule binding experiments with the LUMICKS C-Trap

Single-molecule experiments were performed using a LUMICKS C-Trap instrument. To position DNA substrates for binding analysis, a standard protocol for making DNA tethers was utilized, in which streptavidin coated polystyrene beads were placed in channel one, biotinylated DNA substrates placed in channel two, binding buffer (20 mM HEPES pH 7.5, 10% glycerol, 1 mM DTT, 1 mM MgCl_2_) placed in channel three, and proteins of interest (i.e., CSB and/or SNM1A) placed in channel 4. Valves were opened and each channel flowed at a pressure of 0.3 bar to maintain laminar flow while catching beads in each optimcal trap. The traps were then moved to channel two and distance varied between the traps while looking for a force response to tether a DNA substrate between the two beads as expected by the extensible worm-like chain model, with 48.5 kb for lambda DNA and 12 kb for the LUMICKS handle ligation products. For 12 kb substrates, the trapping laser power set to 15% overall power and 1.5-1.7 μm streptavidin beads were utilized (LUMICKS), and for the lambda DNA substrates, laser power was set to 30% overall power and 4.4-4.8 μm streptavidin beads were used (LUMICKS).

After obtaining single DNA tethers, channels three and four were flowed at 0.3 bar to introduce proteins of interest into the flow cell while keeping the DNA substrate in the buffer alone. DNA tension was then defined in the absence of laminar flow (after testing various tensions for collection, 20 pN at 30°C was determined to be optimal for binding) and then kymographs were collected when the DNA moved to the protein channel.

### Confocal imaging

HaloTag-JF-552 was excited at 561 nm and emission collected in a 575-625 nm band pass filter, and HaloTag-JF-635 was excited with a 638 nm laser and emission collected in a 650-750 nm band pass filter. Data were collected with a 1.2 NA 60X water emersion objective and fluorescence measured with single-photon avalanche photodiode detectors. Each laser was set to 5% power (1-2 µW), pixel size set to 100 nm, scan rates at 30 frames per second.

### Data analysis

Single molecule C-Trap instrument ??OK?? data was exported with Bluelake and analyzed using custom software from LUMICKS (Pylake). Positional and time data for each tracked event was used to determine the dwell time and the duration of gaps between lines for on rate calculation. For each dataset, dwell times were sorted by length to obtain a cumulative residence time distribution plot (CRTD) and fit to exponential decay functions to obtain off rates as previously reported^42^. To determine fitting to a one or two phase decay model, Aikake’s information criteria test was performed after modelling both regimes, with the results given in Table S7 and S8. For two phase exponential decays, for comparison of different datasets, the lifetimes were averaged by summing the lifetimes multiplied by the percentage of total events. To measure on rates, the gaps between the events at the same position were calculated and again fit to an exponential decay function in a cumulative gap time distribution plot (CGTD). This gave apparent on rates which were corrected Concentrations were measured by interpolating the background fluorescence to a standard curve, and correcting the measurement for the presence of free halotag ligand by ratiometrically subtracting the signal as observed by SDS-PAGE quantification. Apparent dissociation constants were determined by correcting the on rate with the protein concentration and using the reciprocal of the average lifetime (from CRTD plot) for determination of the off rate as discussed in Rakowski *et al* ^76^.

### Western blotting

A 10 μL aliquot of the nuclear extracts used for single-molecule studies at 0.93 mg/mL (as determined by Bradford assay) was diluted with 20 μL of water and prepared by boiling at 98°C for 5 min with 1x NuPAGE LDS Sample Buffer (Invitrogen) and 1x NuPAGE Sample Reducing Agent. 20 μL of the prepared nuclear extract was loaded onto a NuPAGE 4-12% Bis-Tris 1.5 mm gel (Invitrogen) and electrophoresis was carried out under a constant voltage of 150 V in 1x NuPAGE MOPS SDS Running Buffer until the dye front reached the end of the gel. Gels were transferred overnight to 0.45 μM pore size Immobilon-P PVDF Membrane (Merck) at 4°C under a constant voltage of 35 V in 1x transfer buffer (25 mM Tris, 192 mM glycine, 20% (v/v) methanol). The following day, membranes were allowed to dry for one hour, then rewetted with 100% methanol. Membranes were then washed 3x 1 min with Tris-buffered saline (TBS), before blocking in 5% milk in TBS for 1 h at room temperature. Membranes were incubated at 4°C overnight with the following primary antibodies in 5% milk in TBS-T (0.1% Tween-20 in TBS): rabbit anti-CSB (Abcam, ab96089, 1:500), rabbit anti-SNM1A (Bethyl, A303-747A, 1:2000), Mouse anti-Pol II (Santa Cruz Biotechnology, CTD4H8, 1:1000), rabbit anti-Histone H3 (Proteintech, 17168-1-AP, 1:5000). Membranes were then washed three times at room temperature for 10 min with TBS-T, before incubation with the following horseradish peroxidase (HRP)-conjugated secondary antibodies, in 5% milk TBS-T for 2 h at room temperature: goat anti-rabbit IgG (H+L) secondary antibody HRP (Invitrogen, 31460, 1:2000) or goat anti-mouse IgG (H+L) secondary antibody HRP (Invitrogen, 31430, 1:2000). Membranes were washed three further times for 10 min each with TBS-T, before membranes were developed using SuperSignal West Pico PLUS Chemiluminescent Substrate (ThermoFisher) and imaged using the iBright 1500 imaging system (Invitrogen).

