## Supplementary Information for "Molecular basis for CSB stimulation of the SNM1A DNA repair nuclease"

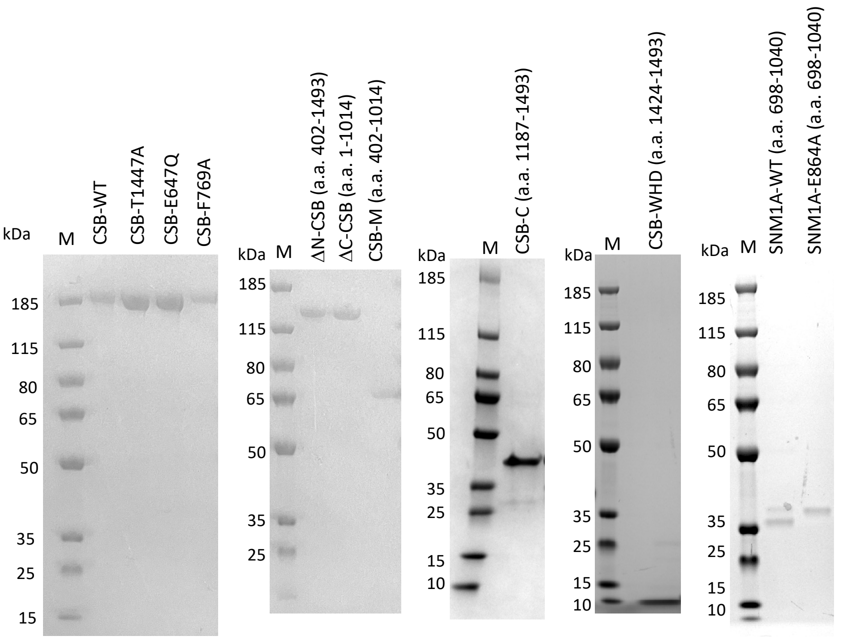

**Figure S1: SDS-PAGE of purified proteins used**. M = Prestained PageRuler Plus protein ladder. Predicted protein masses are: FL-CSB (WT, T1447A, E647Q, F796A) – 180 kDa, ΔC-CSB – 136 kDa, ΔN-CSB – 132 kDa, CSB-M – 72 kDa, CSB-C – 42 kDa, CSB-WHD – 8 kDa, SNM1A (WT + E864A) – 37 (untagged) and 42 kDa (tagged).

**Table S1: Parameters from single-cycle SPR fitting.** Experiments performed where N-Avi-Biotin-ΔN-SNM1A proteins (WT or E864A) were immobilized on the chip. See Methods section for experimental details.

| | | **Immobilized** | | |
|---|---|---|---|---|
| | | **WT-SNM1A** | | **E864A-SNM1A** |
| **Analyte** | **FL-CSB** | $k_a$ = 4.16 x $10^5$ ± 1.1 x $10^4$ $M^{-1}$ $s^{-1}$<br>$k_d$ = 0.001991 ± 4.6 x $10^{-5}$ $s^{-1}$<br>$K_D$ = 4.79 x $10^{-9}$ M<br>Drift = 0.3 RU $s^{-1}$ (fixed)<br>$t_c$ = 1.88 x $10^7$ ± 6.5 x $10^4$ | | Not determined |
| | **CSB (a.a. 1187-1493)** | $k_a$ = 3.82 x $10^5$ ± 6.3 x $10^3$ $M^{-1}$ $s^{-1}$<br>$k_d$ = 0.004048 ± 6.5 x $10^{-5}$ $s^{-1}$<br>$K_D$ = 1.06 x $10^{-8}$ M<br>Drift = 0.3894 RU $s^{-1}$ (fitted)<br>$t_c$ = 3.01 x $10^7$ ± 9.2 x $10^4$ | $k_a$ = 5.61 x $10^5$ ± 3.3 x $10^2$ $M^{-1}$ $s^{-1}$<br>$k_d$ = 0.007121 ± 4.2 x $10^{-6}$ $s^{-1}$<br>$K_D$ = 1.27 x $10^{-8}$ M<br>Drift = 0.3894 RU $s^{-1}$ (fitted)<br>$t_c$ = 4.07 x $10^7$ ± 1.9 x $10^4$ | $k_a$ = 3.76 x $10^5$ ± 1.0 x $10^4$ $M^{-1}$ $s^{-1}$<br>$k_d$ = 0.01098 ± 3.0 x $10^{-4}$ $s^{-1}$<br>$K_D$ = 2.92 x $10^8$ M<br>Drift = 0.845 RU $s^{-1}$ (fitted)<br>$t_c$ = 4.54 x $10^7$ ± 5.9 x $10^5$ |
| | **CSB (a.a. 1424-1493)** | $k_a$ = 1.02 x $10^5$ ± 3.0 x $10^3$ $M^{-1}$ $s^{-1}$<br>$k_d$ = 0.009041 ± 2.5 x $10^{-4}$ $s^{-1}$<br>$K_D$ = 8.90 x $10^8$ M<br>Drift = 0.1231_RU $s^{-1}$ (fitted)<br>$t_c$ = 3.12 x $10^6$ ± 1.3 x $10^4$ | | $k_a$ = 644.7 ± 1.1<br>$k_d$ = 7.04 x $10^{-4}$ ± 2.9 x $10^{-6}$ $s^{-1}$<br>$K_D$ = 1.09 x $10^6$ M<br>Drift = 0.06 RU $s^{-1}$ (fixed)<br>$t_c$ = 3.00 x $10^6$ (fixed) |

**Table S2: Parameters from single-cycle SPR fitting when wild-type N-His-ZB-CSB-Avi-Biotin was immobilized on the chip.** See Methods section for experimental details.

| Analyte | WT-SNM1A | E864A-SNM1A |
|---|---|---|
| Parameters determined | $k_a = 6.88 \times 10^5 \pm 5.8 \times 10^3\ M^{-1}\ s^{-1}$<br>$k_d = 0.002993 \pm 1.8 \times 10^{-5}\ s^{-1}$<br>$K_D = 4.35 \times 10^{-9}$ M<br>Drift = 0.3 RU $s^{-1}$ (fixed)<br>$t_c = 6.80 \times 10^7$ | $k_a = 1.71 \times 10^5 \pm 1.1 \times 10^3\ M^{-1}\ s^{-1}$<br>$k_d = 0.001386 \pm 5.7 \times 10^{-6}\ s^{-1}$<br>$K_D = 8.11 \times 10^{-9}$ M<br>Drift = 0.3 RU $s^{-1}$ (fixed)<br>$t_c = 6.80 \times 10^7$ |

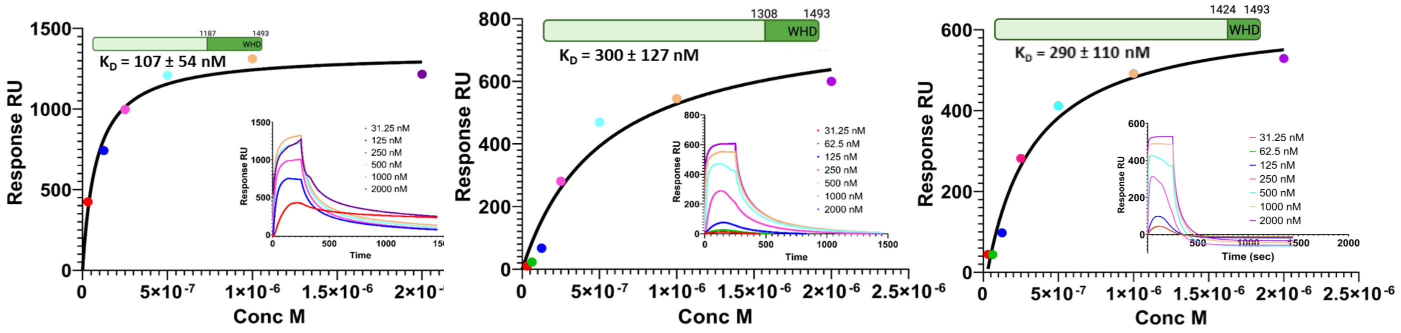

**Figure S2: Multi-cycle and dose response curves for CSB (a.a. 1187-1493), CSB (a.a. 1308-1493) and CSB (a.a. 1424-1493).** N-Avi-Biotin-SNM1A was immobilised on the chip. Dissociation constants are given. For CSB a.a 1187-1493, data for analyte at 62.5 nM was omitted due to the presence of an air bubble during the injection.

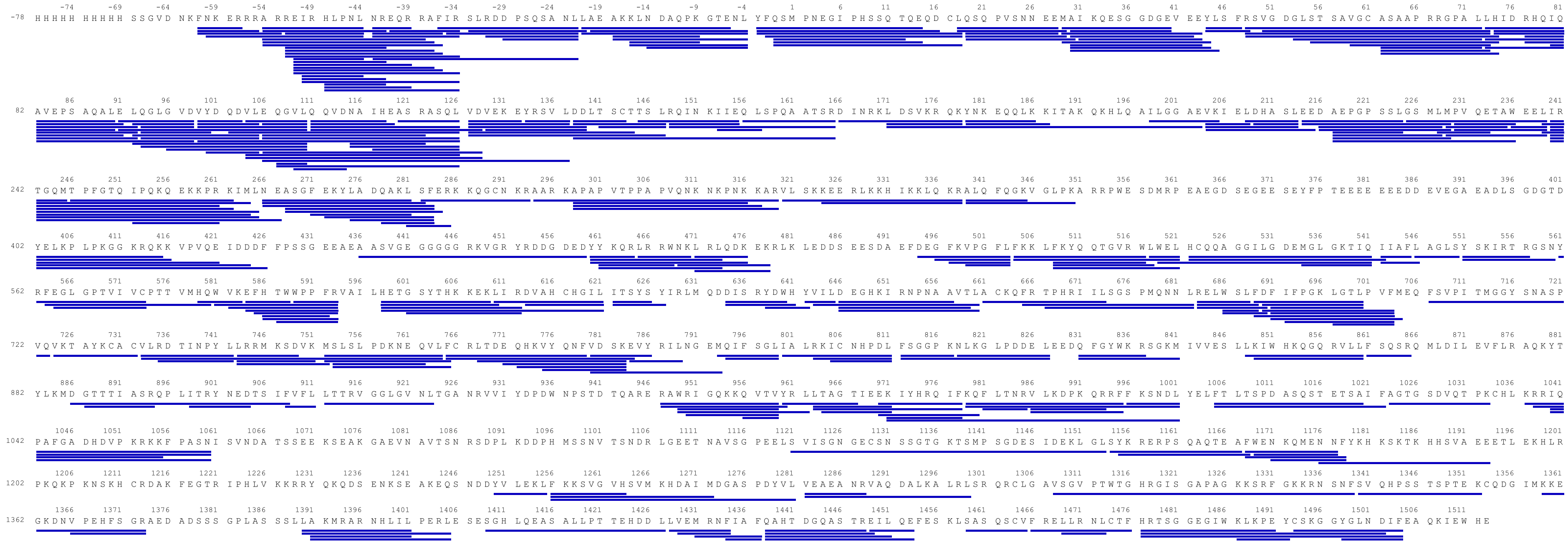

**Fig S3: Peptide coverage obtained for CSB in HDX-MS experiments.** 376 peptides were observed with 80% sequence coverage, 3.1 average redundancy.

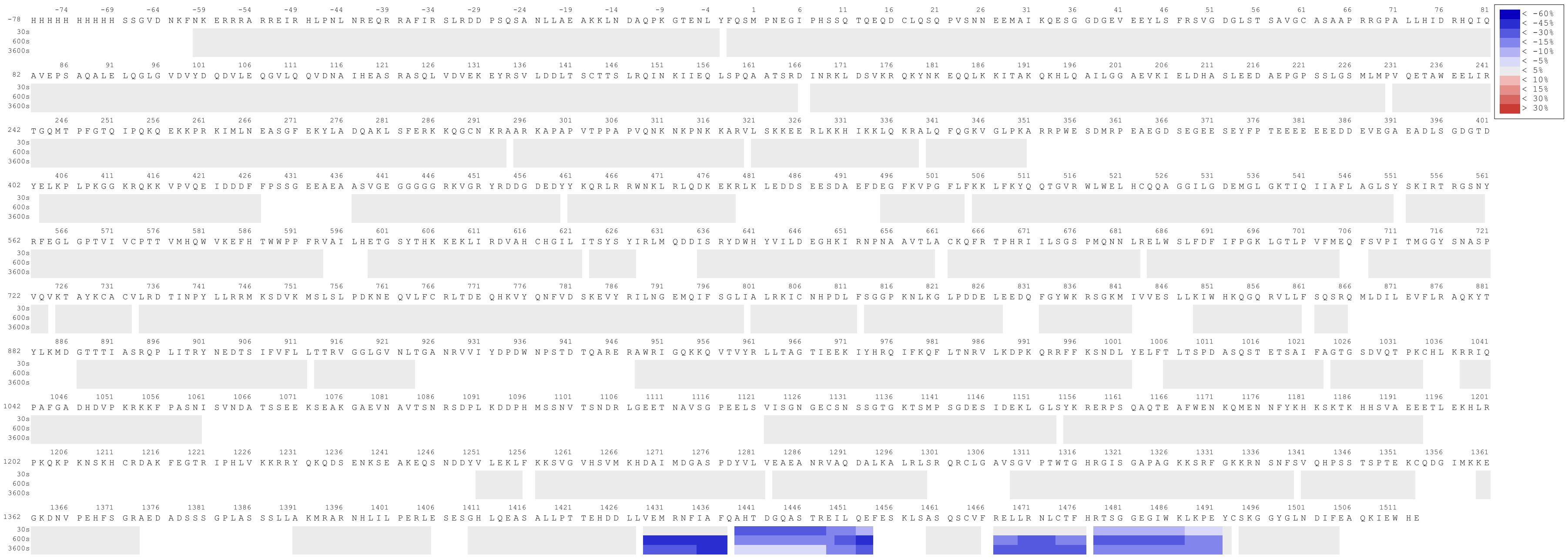

**Figure S4: CSB peptides with changes in deuterium uptake in HDX-MS experiments upon interaction with SNM1A.**

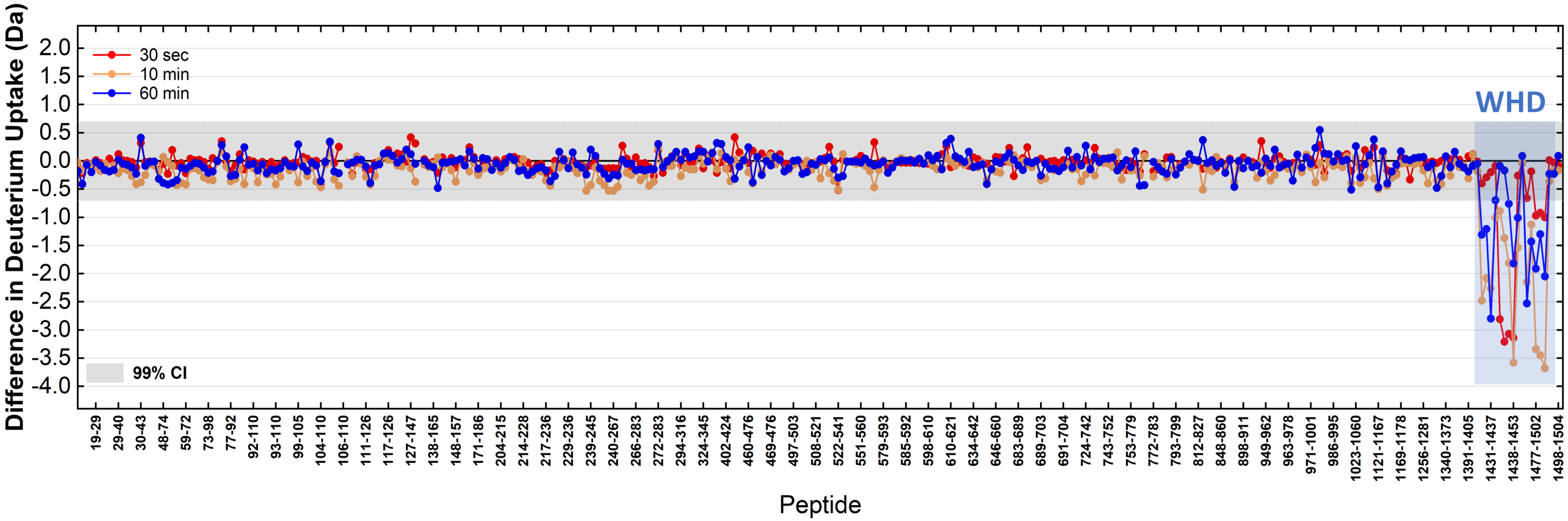

**Fig S5: Changes in deuterium uptake in the CSB protein upon interaction with SNM1A are localised to the CSB-WHD.**

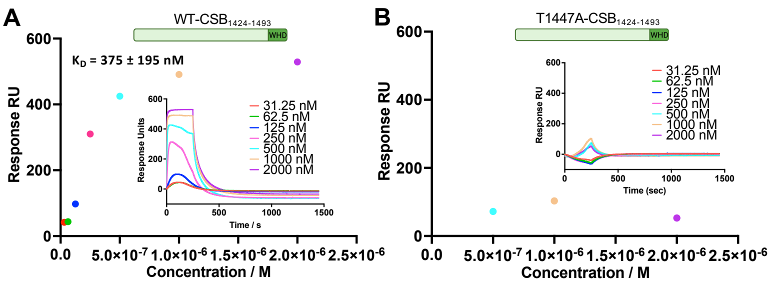

**Fig S6: Multi-cycle and dose response curves for CSB (a.a. 1424-1493, wild-type T1447A.** N-Avi-Biotin-SNM1A was immobilised on the chip. Dissociation constants are given.

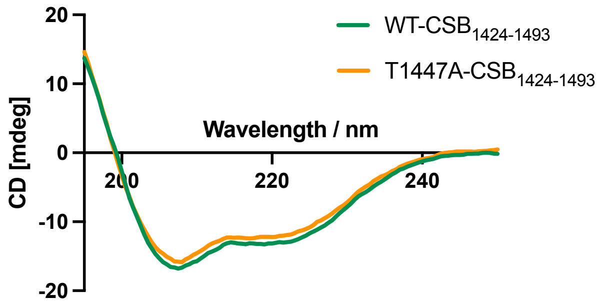

**Fig S7: Circular dichroism spectroscopy of isolated CSB-WHD**. Wild-type and T1447A.

**Table S3: Oligonucleotide sequences used for biochemical assays.** P refers to monophosphate.

| Code | Sequence of oligonucleotide (5' to 3') |
|---|---|
| WN1 | P-CAATTAGCTGAGGGTCACGTTTAATTAATTATTTACGGAAGAGTGTGAAG-Biotin |
| WN10 | Biotin-TCTTCACACTCTTCCGTTAATAATTAATTAAACGTGACCCTCACGTAATT |
| WN9 | P-AACGGAAGAGTGTGAAG |
| WN2 | Biotin-TCTTCACACTCTTCCGTAGGACCGACGGATTTGGTGACCCTCAGCTAATT |
| WN10 short | Biotin-TCTTCACACTCTTCCGTTAATAATT |
| '0-nt' ICL | Top strand: **Y**ACGGAAGAGTGTGAAG (Y = azide)<br>Bottom strand: TCTTCACACTCTTCCGT**X**GGACCGACGGATTTGGTGACCCTCAGCTAATT (X = alkyne) |

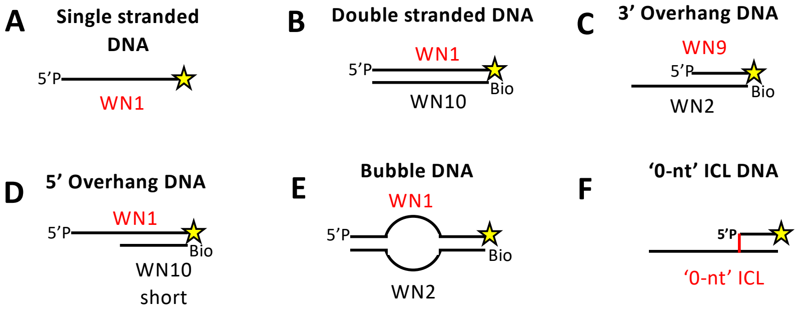

**Figure S8: DNA structures used in biochemical assays.** The annealing protocol is described in the Methods section. 5'P denotes a 5' phosphate and Bio represents a biotin modification. The radiolabeled oligonucleotide is given in red, and the location of labelling annotated by a yellow star.

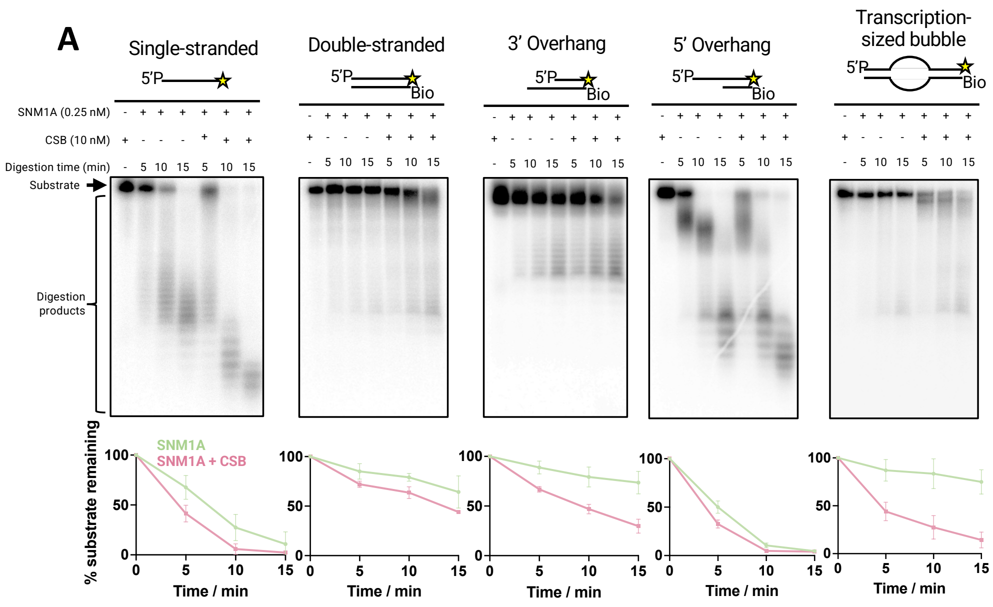

**Figure S9: The substrate scope of CSB stimulation of SNM1A activity.** Oligonucleotides used (with preparation method) are given in Table S3 and Fig. S8. The location of the 3′ 32P is given by a yellow star, Bio represents the location of 5′ biotins on the oligo and 5′P the location of a 5′ phosphate group. Quantification was based upon assays conducted in triplicate; error bars represent one standard deviation. Time course assays of SNM1A activity on a panel of DNA repair intermediate substrates in the absence of WT-CSB.

**Table S4: Oligonucleotides used to prepare ICL-containing DNA repair intermediate for single-molecule studies.**

| | |
|---|---|
| Top strand 1 | CAAC GCTCTTC AATTAGCTGAGGGTCAC |
| Top strand 2 | P-GTTTAATTAATTATT **Y** ACGGAAGAGTGTGAAG (Y = azide) |
| Bottom strand | ACCA CTTCACACTCTTCCGT **X** GGACCGACGGATTTG GTGACCCTCAGCTAATT GAAGAGC (X = alkyne) |

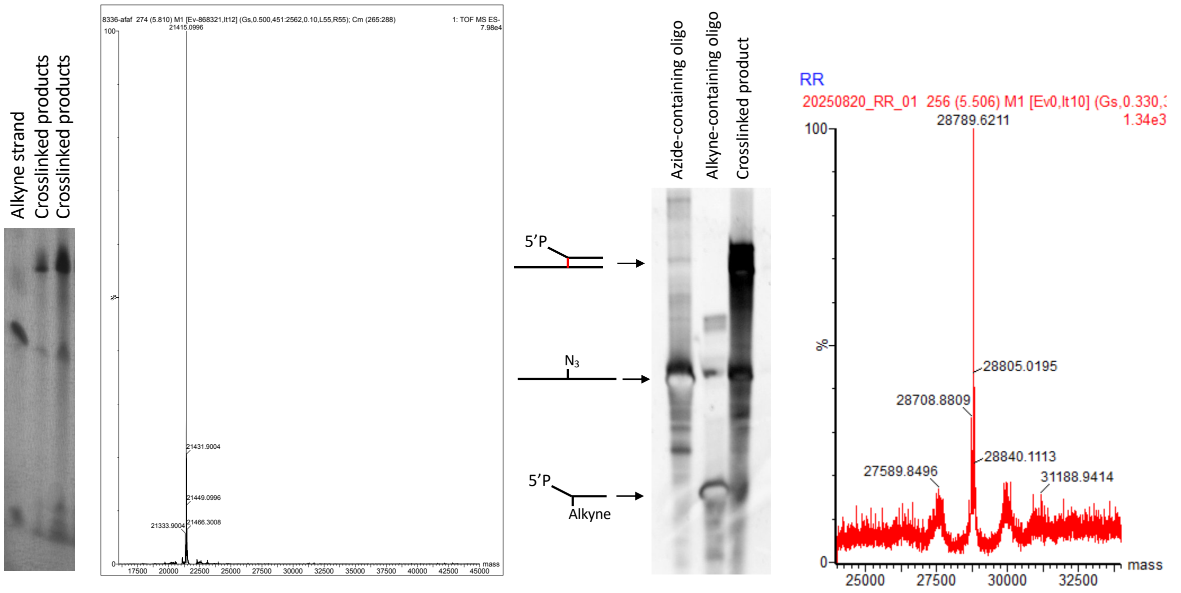

**Figure S10: Isolation of Cu(I)-click ICL-containing oligonucleotides. A.** Gel purification of the '0-nucleotide' ICL substrate used for biochemical assays. 8% denaturing PAGE gel showing the un-reacted alkyne-containing strand (left lane) and the reaction products from the Click-iT reaction between the azide- and alkyne-containing strands (all other lanes). The larger, 'clicked' products were excised from the gel and purified as described in the Methods section. See Supplementary Figure 1 for oligonucleotide structure. **B.** Q-TOF MS of the '0-nt ICL' substrate. Expected [M+H]- *m/z* = 21414.5, observed [M+H]- m/z = 21415.0996 Da. **C.** 10% denaturing PAGE gel of the ICL-containing substrate used for single-molecule studies, showing the azide and alkyne-containing oligos alone, and post crosslinking. The crosslinked product was run out on a 10% denaturing PAGE gel and the band corresponding to the higher molecular weight product was excised from the gel and purified as detailed in the Methods section. **D.** Q-TOF MS of the ICL-containing oligonucleotide tethered in the LUMICKS' C-Trap. Expected [M+H]- *m/z* = 28789.1, observed [M+H]- *m/z* = 28789.6211.

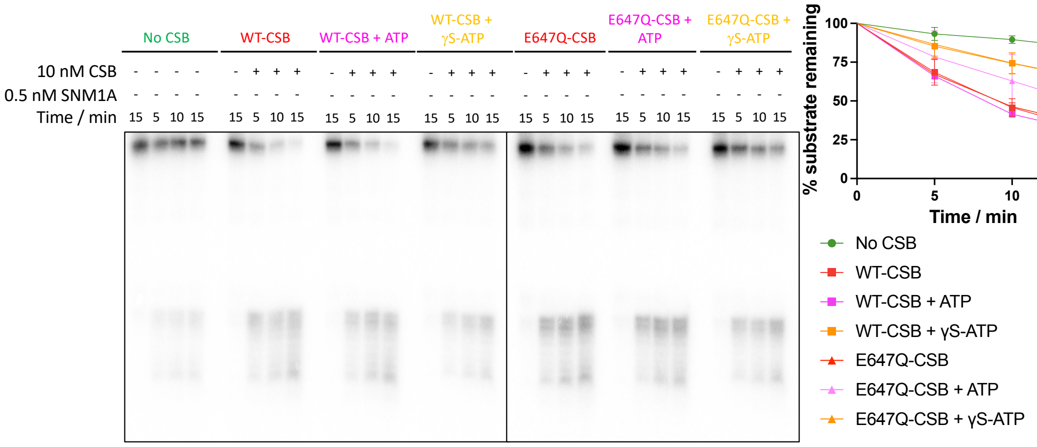

**Figure S11: Evidence that the ATPase activity of CSB is not involved in the stimulation of SNM1A activity**. The presence of ATP does not affect the enhancement of SNM1A activity by CSB. ATP was used at 50 μM final and hydrolysis observed on the '0-nucleotide ICL' substrate used.

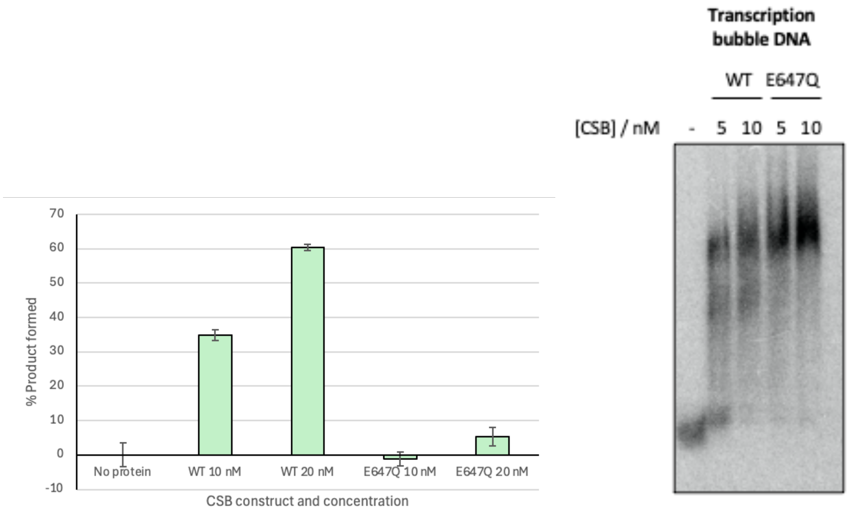

**Figure S12: E647Q-CSB is ATPase-dead, and this does not affect the DNA binding ability of CSB.** ATPase reactions were performed as detailed in the methods section using the Malachite Green assay, detecting for release of orthophosphate. EMSA of WT and E647Q.

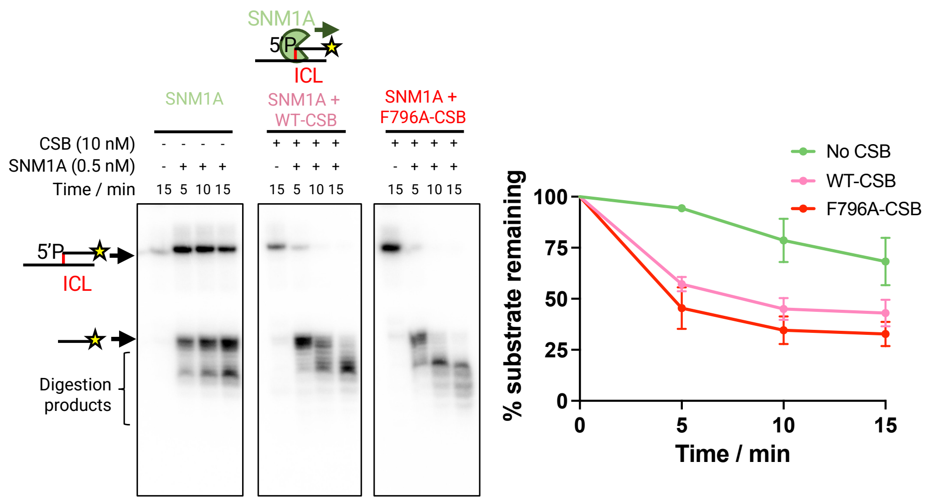

**Figure S13: Pulling hook dead CSB (F796A) is not involved in the stimulation of SNM1A activity**

.

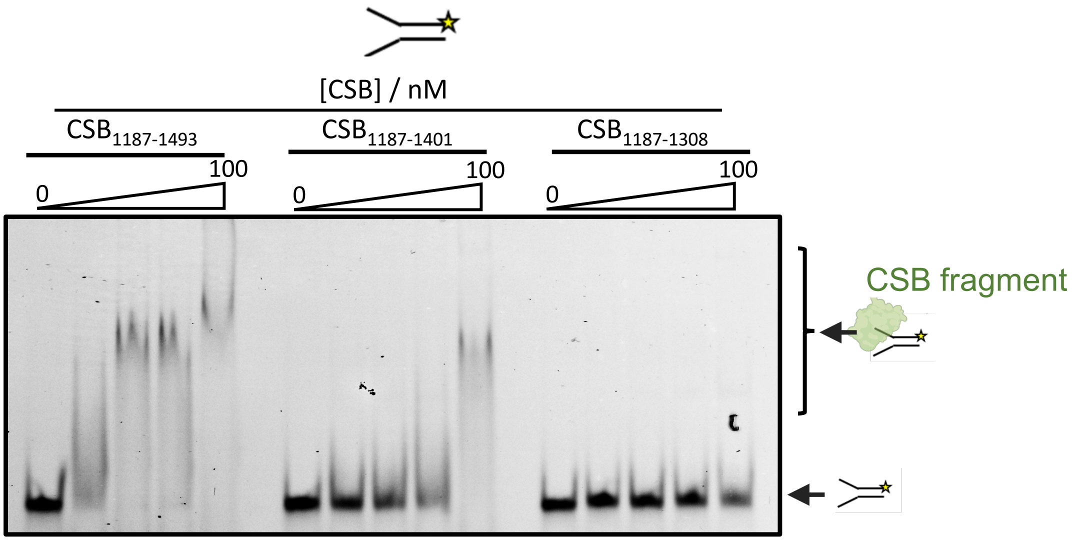

**Figure S14: EMSA of C-terminal domains of CSB binding to DNA.** Fork DNA (as reported in Abdullah *et al*. 2017) was chosen as the substrate because it possesses both double stranded and single stranded character. CSB fragments $CSB_{1308-1493}$ and $CSB_{1424-1493}$ were not found to bind DNA (data not shown). The results are representative of three repeats.

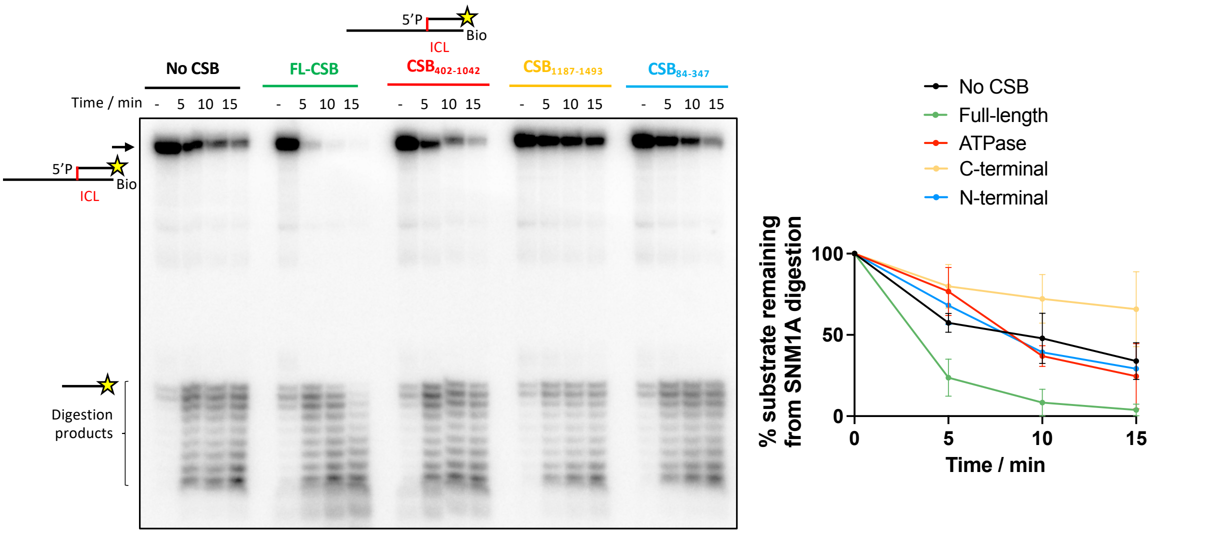

**Figure S15: Nuclease assay of SNM1A activity in the presence of different CSB domains.** Quantification is performed from three repeats with the standard deviation given in the error bars.

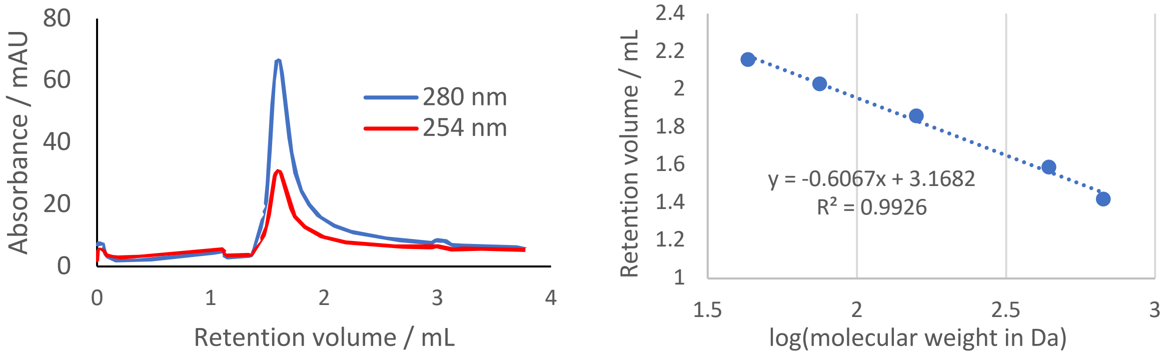

**Figure S16: Akta trace of purified full-length CSB with corresponding calibration curve**. The peak corresponds to a retention volume of 1.56 mL on Superdex S200 GL 5/150, corresponding to a molecular weight of 440 kDa (predicted dimer = 336 kDa) based on the calibration curve given. The standards used were thyroglobulin (669 kDa), ferritin (440 kDa), aldolase (158 kDa), conalbumin (75 kDa) and ovalbumin (43 kDa).

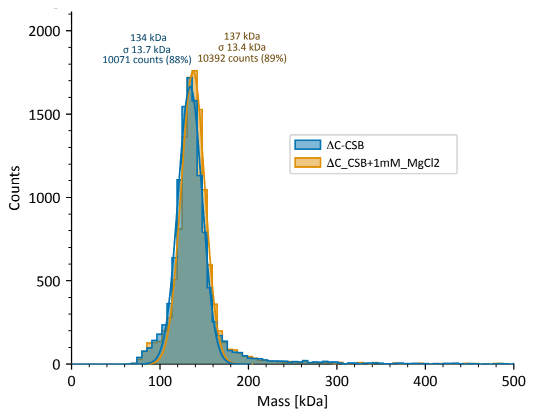

**Figure S17: Mass photometry of ΔC-CSB and ΔN-CSB. A.** MP of ΔC-CSB. Expected monomer = 136 kDa, expected dimer = 272 kDa. **B.** MP of ΔN-CSB. Expected monomer = 132 kDa, expected dimer – 264 kDa. Measurements were performed in the absence/presence of $MgCl_2$ and the results imply that $Mg^{2+}$ does not influence CSB dimerisation. Experiments were performed as described in Materials and Methods.

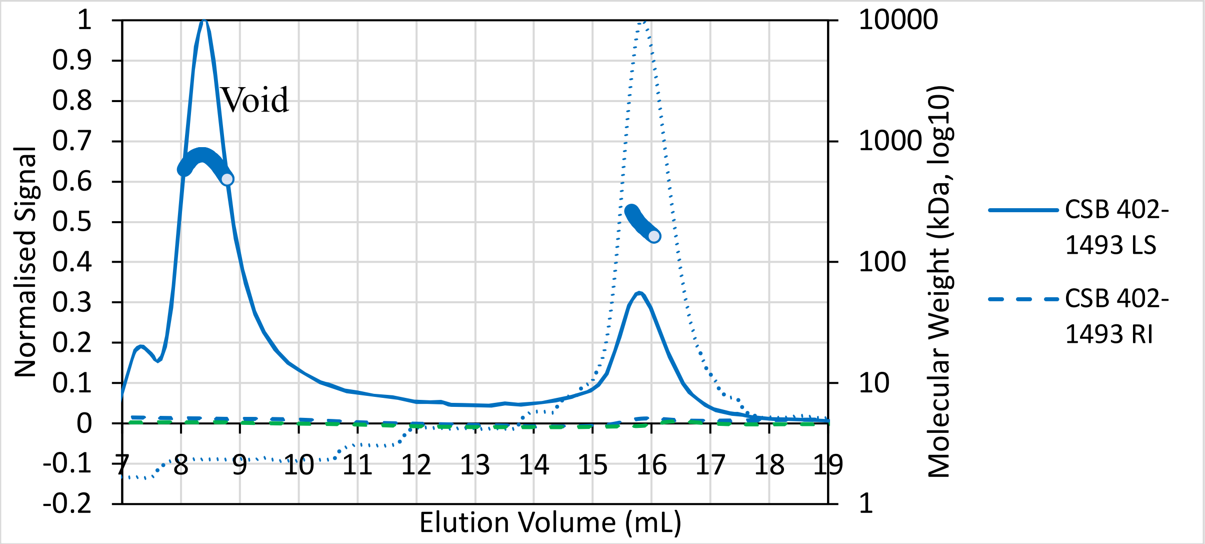

**Figure S18: SEC-MALs of ΔN-CSB**. The peak at 15.8 mL corresponds to ΔN-CSB of molecular weight 199 kDa. Expected monomer = 132 kDa, expected dimer – 264 kDa.

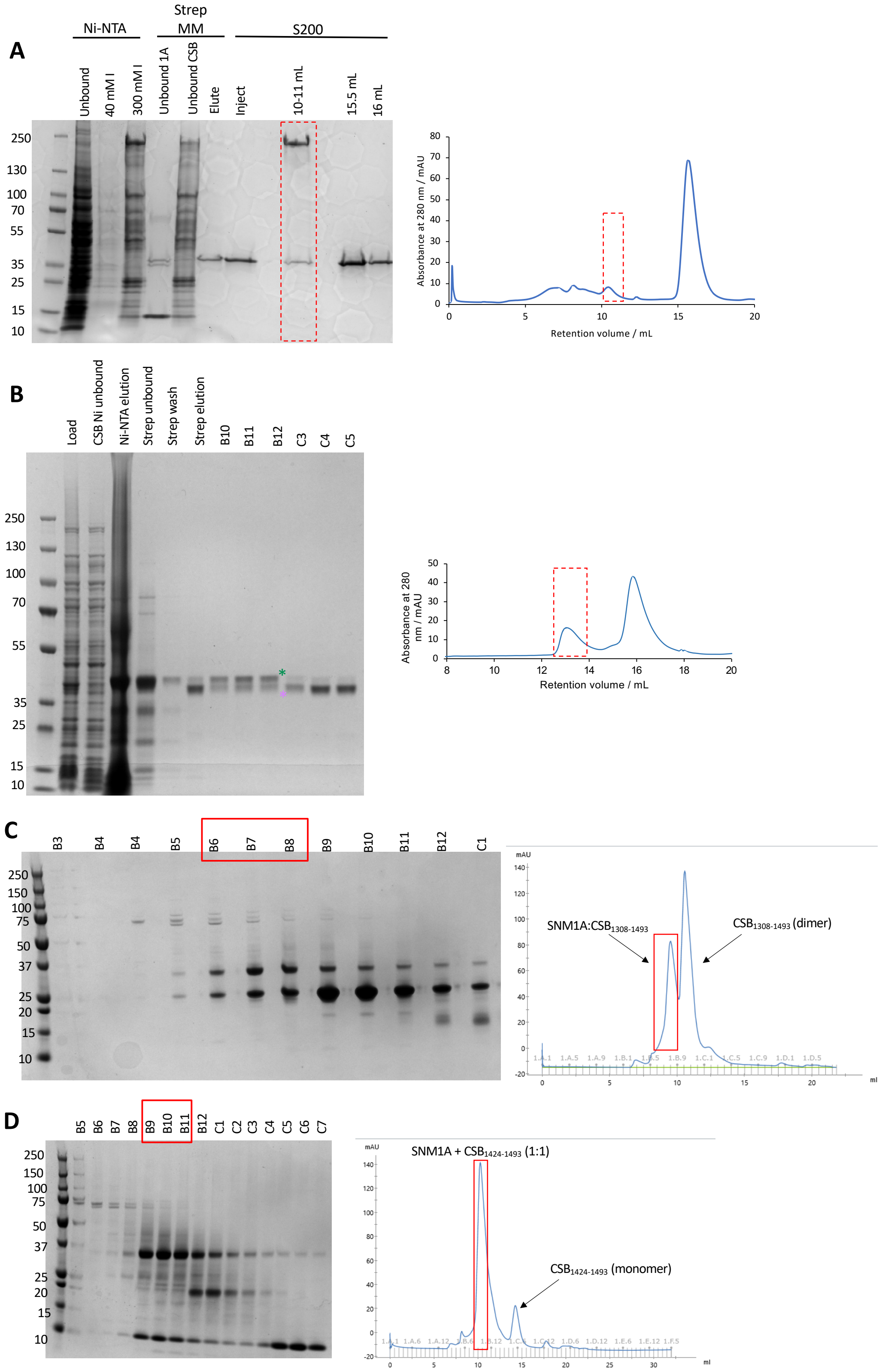

**Figure S19: Co-purifications of CSB and SNM1A.** Methods for purifying each complex are detailed in the Methods section. Figures shown from size-exclusion chromatography traces of eluted protein complexes on Superdex S75 GL 10/300 or Superdex S200 GL 10/300.

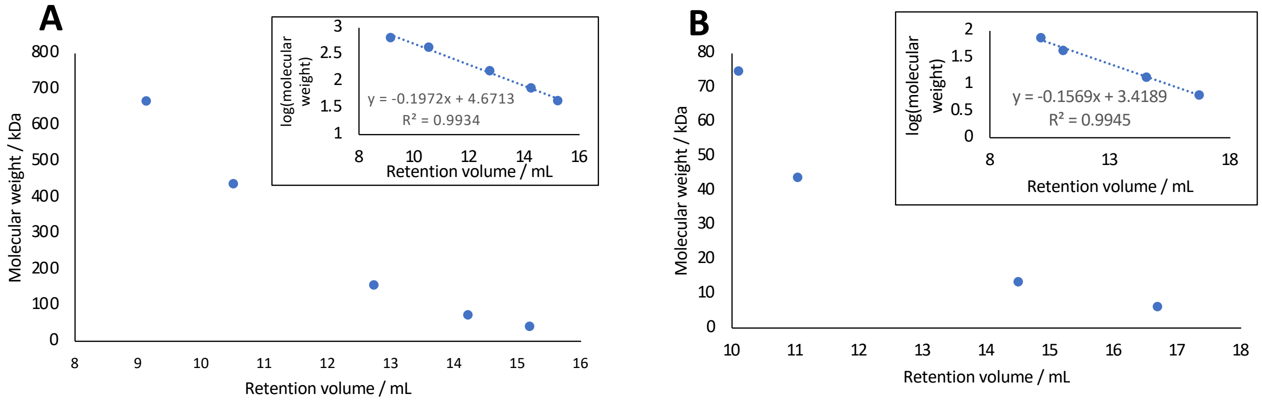

**Figure S20: Size exclusion calibration curves. A.** Superdex S200 GL 10/300. The standards used were thyroglobulin (669 kDa), ferritin (440 kDa), aldolase (158 kDa), conalbumin (75 kDa) and ovalbumin (43 kDa). **B.** Superdex S75 GL 10/300. The standards used were conalbumin (75 kDa), ovalbumin (43 kDa), ribonuclease A (13.7 kDa), aprotinin (6.5 kDa).

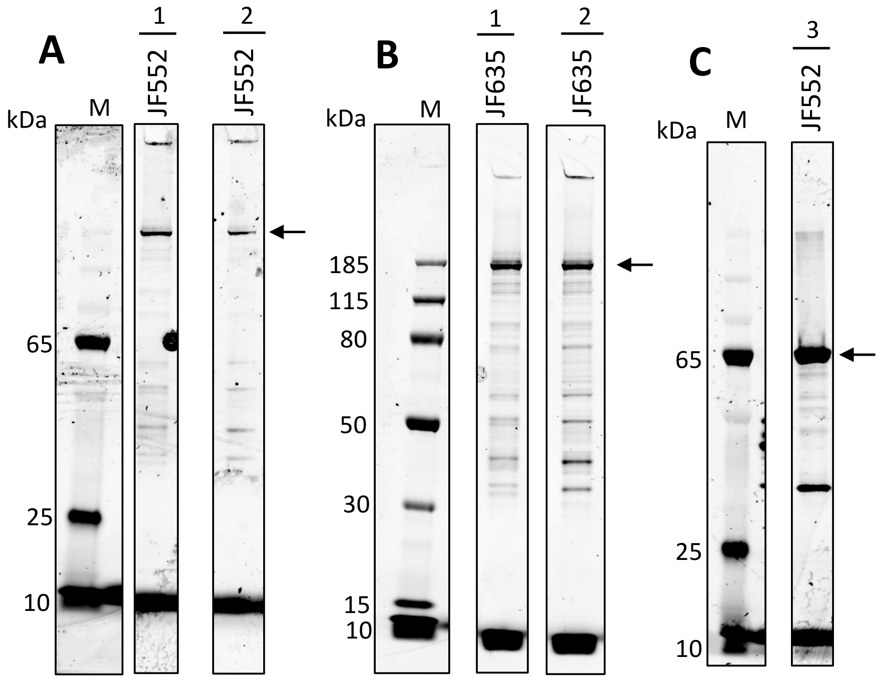

**Figure S21: SDS-PAGE gels of nuclear extracts used for single-molecule studies**. A. Halotag-CSB constructs labelled with JF552, imaged using a Typhoon imager at 532 nm. B. Halotag-CSB constructs labelled with JF635, imaged using a Typhoon imager at 647 nm. C. Partially purified Halo-SNM1A (a.a. 698-1040) labelled with JF552, imaged using a Typhoon imager at 532 nm.

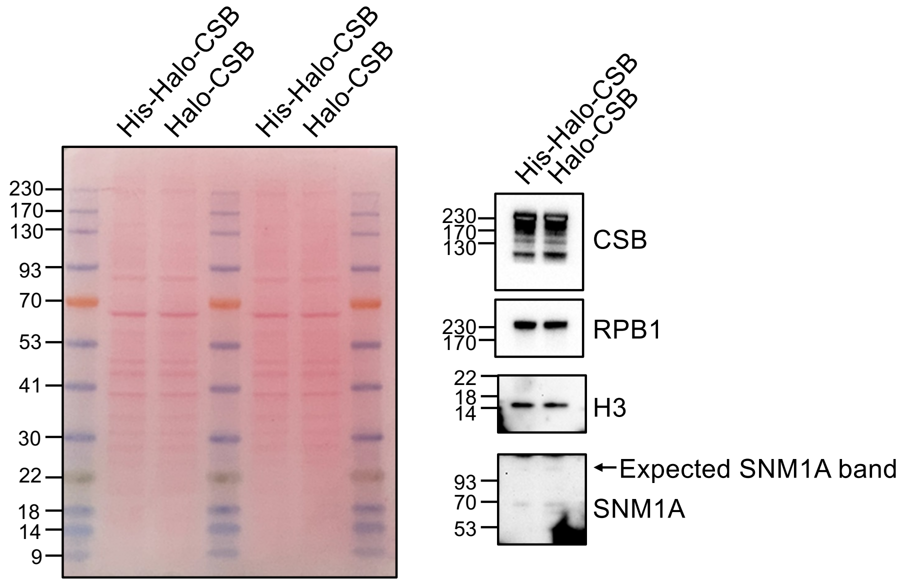

**Figure S22: Western blot analysis of CSB-transfected nuclear extracts**. Ponceau staining of membrane with western blotting for CSB, the major RNAPII subunit, RPB1, H3 and SNM1A.

**Table S5: Parameters determined for protein:DNA binding events on lambda DNA in the LUMICKS' C-Trap.**

| Dataset | Final protein concentration (nM) | Recruitment time of SNM1A to bound CSB (s) | Apparent $k_{on}$ ($s^{-1}$) | Corrected $k_{on}$ ($M^{-1} s^{-1}$) | Lifetime(s) and percentages (s) | Binding lifetime $t_{avg}$ (weighted average, s) | Weighted $k_{off}$ ($s^{-1}$) | $K_D$ (nM) | Oligomerisation state |
|---|---|---|---|---|---|---|---|---|---|
| SNM1A | 2.23 ± 0.426 | - | 0.0403 ± 0.0405 | 1.81 ± 1.81 × $10^7$ | 8.64 ± 0.548 (23 ± 0.6%)<br>0.706 ± 0.0107 (77 ± 0.6%) | 2.53 | 0.395 | 21.8 ± 10.9 | Monomer |
| CSB | 0.547 ± 0.0351 | - | 0.0156 ± 0.00882 | 2.86 ± 1.05 × $10^7$ | 73.2 ± 24.1 (34 ± 5%)<br>7.12 ± 0.926 (66 ± 5%) | 29.6 | 0.0338 | 1.18 ± 0.667 | CSB 73:27 monomer:dimer |
| Co-localised SNM1A + CSB | SNM1A: 2.23 ± 0.426<br>CSB: 2.38 ± 0.268 | 71.9 ± 5.52 (79 ± 3%)<br>3.56 ± 0.95 (21 ± 3%)<br>Average: 57.5 ± 1.42 s | 0.017 ± 0.000420 | 7.80 ± 0.72 × $10^6$ | 16.2 ± 2.52 (42 ± 2%)<br>0.825 ± 0.0955 (58 ± 2%) | 7.28 | 0.137 | 17.5 ± 1.62 | CSB 27:73 monomer:dimer |

**Table S6: Parameters determined for protein:DNA binding events on the ICL lesion in the LUMICKS' C-Trap.**

| Dataset | Final protein concentration (nM) | Recruitment time of SNM1A to bound CSB (s) | Apparent $k_{on}$ ($s^{-1}$) | Corrected $k_{on}$ ($M^{-1} s^{-1}$) | Lifetime(s) and percentages (s) | Binding lifetime $t_{avg}$ (weighted average, s) | Weighted $k_{off}$ ($s^{-1}$) | $K_D$ (nM) | Oligomerisation state |
|---|---|---|---|---|---|---|---|---|---|
| SNM1A | 2.99 ± 0.626 | - | 9.36 ± 7.57 × $10^{-3}$ | 3.13 ± 3.19 × $10^6$ | 0.901 ± 3.19 | 0.901 | 1.11 | 355 ± 361 | Monomer |
| Co-localised SNM1A + CSB | SNM1A: 13.2 ± 1.13<br>CSB: 2.95 ± 0.463 | 40.6 ± 7.3 s | 0.0246 ± 0.00372 | 1.87 ± 0.16 × $10^6$ | 31.1 ± 17.9 (33 ± 12%)<br>2.26 ± 0.783 (67 ± 12%) | 11.8 | 0.0847 | 45.3 ± 3.88 | CSB 8:92 monomer:dimer |

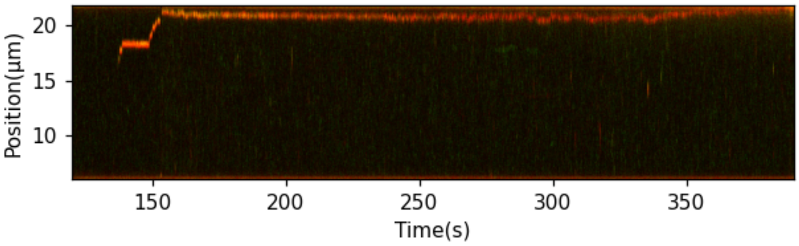

**Figure S23: Kymograph showing 2 different colours of CSB protein diffusing together on lambda DNA.**

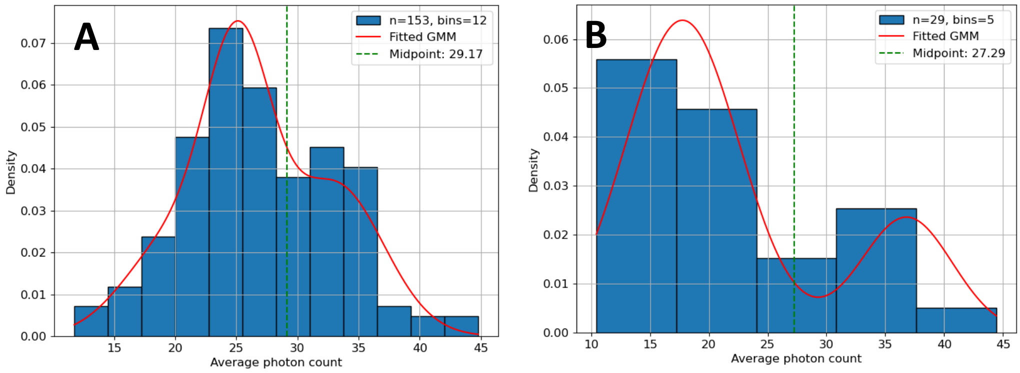

**Figure S24: Histograms of average CSB photon counts along a kymotrack**. To determine whether a CSB:DNA binding event was with a monomeric or dimeric CSB, the average photon count across an entire kymotrack was determined by taking the average of a line of best fit across the plotted photon count intensities for the track. The values for each CSB binding event were combined and shown as histograms. **A.** CSB average photon counts on lambda DNA. Events with an average photon count >29.17 were denoted as dimers, and <29.17, monomeric. **B.** CSB average photon counts for binding events localised at the ICL. Events with an average photon count >27.29 were denoted as dimers, and <27.29, monomeric.

**Table S7: Statistical analysis (AICc test) to determine whether binding event lifetimes (CTRD plots) exhibited one- or two-phase decay.** Highlighted in grey are the most negative values i.e. the most appropriate fit for the data.

| DNA substrate | Protein | AICc for one phase decay | AICc for two phase decay |
|---|---|---|---|
| Lambda | SNM1A | -5180 | -6973 |
| Lambda | CSB | -399.9 | -474.9 |
| Lambda | SNM1A + CSB | -550.8 | -767 |
| ICL | SNM1A | -165.6 | -164.3 |
| ICL | SNM1A + CSB | -122.9 | -137.8 |

**Table S8: Statistical analysis (AICc test) of SNM1A recruitment times to DNA-bound CSB**. Highlighted in grey are the most negative values i.e. the most appropriate fit for the data.

| DNA substrate | Protein | AICc for one phase decay | AICc for two phase decay |
|---|---|---|---|
| Lambda | SNM1A + CSB | -612.4 | -699.8 |
| ICL | SNM1A + CSB | -107.2 | -100.4 |
